# Single Nucleotide Resolution 4sU Sequencing (SNU-Seq) reveals the transcriptional responsiveness of an epigenetically primed human genome

**DOI:** 10.1101/2021.07.14.452379

**Authors:** Umut Gerlevik, Philipp Lorenz, Anna Lamstaes, Harry Fischl, Shidong Xi, Aksel Saukko-Paavola, Struan Murray, Thomas Brown, Alexander Welch, Charlotte George, Andrew Angel, Andre Furger, Jane Mellor

## Abstract

Genomes are pervasively transcribed, leading to stable and unstable transcripts that influence 3-dimensional genome organisation and gene regulation. High sensitivity and nucleotide resolution are required to resolve mammalian nascent transcriptomes. Here, we exploit the sensitivity of 4-thio-uridine (4sU) metabolic pulse-labelling to develop two nucleotide-resolution methods: <u>S</u>ingle-<u>N</u>ucleotide resolution 4s<u>U</u> sequencing (SNU-Seq) and <u>s</u>ize-fractionated 4sU-Seq (sf4sU-Seq). sf4sU-Seq involves gel isolation of abundant 4sU-labelled promoter proximal nascent transcripts, enabling nucleotide resolution mapping of transcription start sites and promoter proximal pauses (PPPs) on the same transcript. SNU-Seq exploits 3’ end RNA-Seq, using bacterial poly(A) polymerase (bPAP) to polyadenylate the 3’ ends of nascent transcripts and create oligo(dT)-primed libraries. The artificial poly(A) tail marks the precise position of polymerase on a transcription unit. SNU-Seq read levels are similar at pre-mRNAs and enhancers genome-wide and read spikes in pre-mRNA outputs map pauses, PPPs and polyadenylation sites. SNU-Seq enables discovery of thousands of unannotated regions of divergent transcription and helps define hundreds of the more than 10,000 regions of primed non-transcribed acetylated open chromatin that induce divergent nascent transcripts within 0.5h of IFN-γ treatment in Hep3B cells. Thus, combining chromatin analysis with SNU-Seq reveals the transcriptional responsiveness of an epigenetically primed human genome.

**HIGHLIGHTS:**

- SNU-Seq maps nascent transcripts with bp resolution, high sensitivity and low cost
- sf4sU-Seq resolves TSS and PPP at the same gene, complementing SNU-Seq
- 1000’s of divergently transcribed enhancers resolved by SNU-Seq
- Rapid IFNγ dependent transcriptional induction from primed Hep3B epigenome

**GRAPHICAL ABSTRACT:** 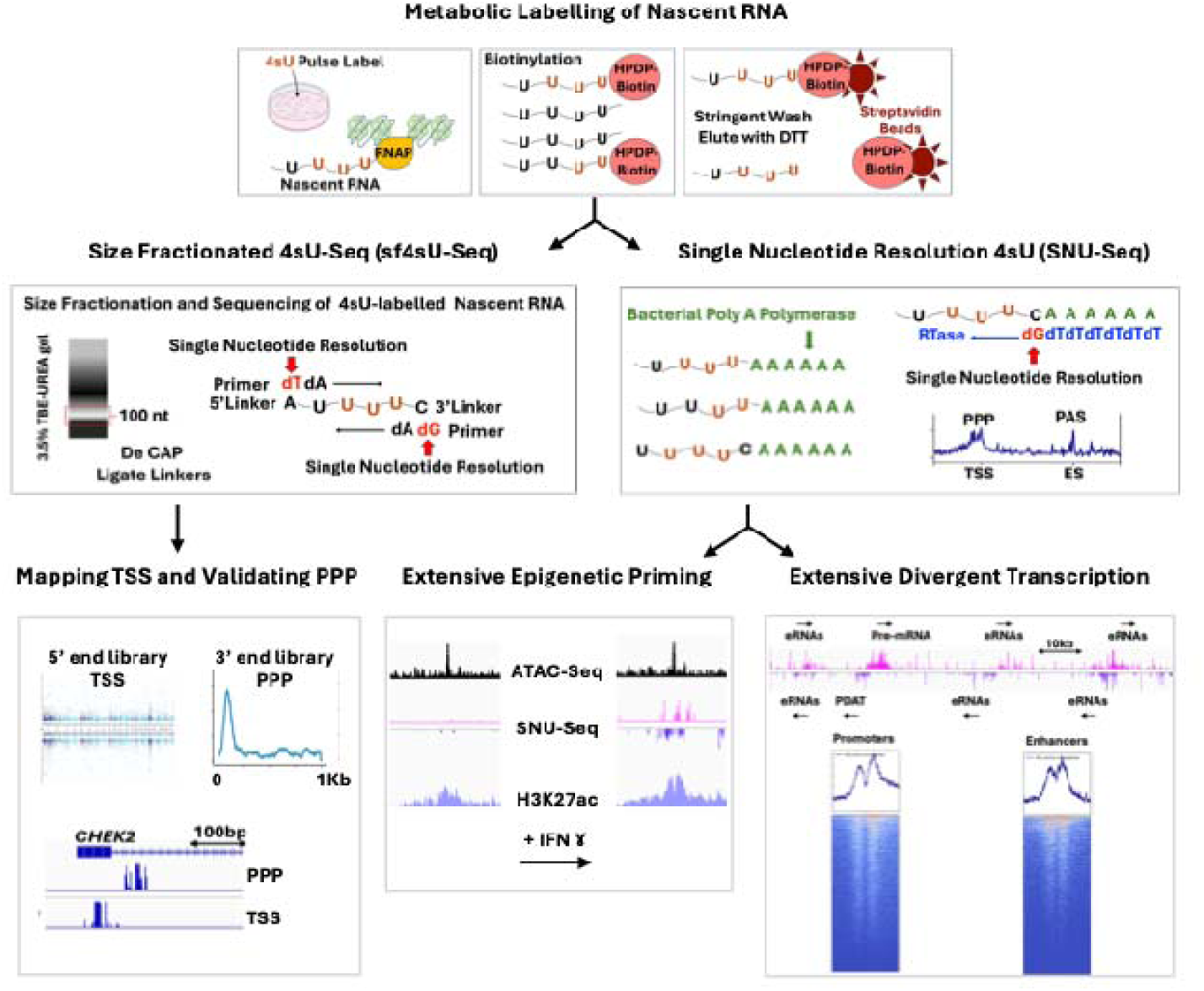

## INTRODUCTION

Genomes are pervasively transcribed, leading to stable and unstable transcripts that influence 3-dimensional genome organisation, the epigenome and gene regulation. Nascent transcription remodels local and higher-order chromatin structures, simply through the act of transcription, through retained nascent transcripts or by recruiting functional RNA-binding proteins, potentiating new combinatorial responses to environmental signals. Routine mapping of precisely where and when such transcription occurs and how it relates to epigenomic features and production of pre-mRNAs requires a low-cost, sensitive method with base-pair resolution to capture co-transcriptional events such as the promoter proximal pause (PPP), pausing at exon-intron boundaries and pauses prior to nascent polyadenylation. Nascent transcripts can be captured using four different approaches: enrichment of nascent transcripts associated with RNA polymerase II (e.g., mNET-Seq) (1–5); enrichment of chromatin-associated nascent transcripts from the nucleus (e.g., ChrRNA-Seq) (6,7); “run-on” approaches in permeabilised cells (e.g., PRO-Seq) (8–12); metabolic pulse-labelling coupled with affinity purification of nascent transcripts (e.g., 4sU-Seq, TT- Seq) (13–19). These methods produce distinct metagene profiles over genes, consistent with each technique capturing different aspects of transcription, but can be challenging to conduct and/or lack base pair resolution (20,21).

Pulse labelling nascent transcripts with 4-thio-uridine (4sU) (22), as in 4sU-Seq (19) and TT- Seq (15), is highly sensitive, as 4sU is rapidly taken up by cells, phosphorylated and incorporated into transcripts. Both 4sU-Seq and TT-Seq provide information about where transcription occurs during the labelling window (usually about 8-10 mins), but transcript processing, library preparation and data smoothing reduce resolution to about 200 nt (TT- Seq) or introduce a 5’ bias (4sU-Seq) in read distribution. Thus, key events in transcription, such as the PPP, are not resolved in TT-Seq metagenes (15,23). Ideally, information about the position of the 3’ end of the intact 4sU-labelled nascent transcripts would provide nucleotide resolution data, as in mNET-Seq or PRO-Seq. To achieve this, <u>S</u>ingle-<u>N</u>ucleotide resolution 4s<u>U</u>-sequencing (SNU-Seq) was developed for use in adherent mammalian cells (HEK293 and Hep3B) with controls so that the precise source of reads can be determined. Like TT-Seq and 4sU-Seq, SNU-Seq takes advantage of a 4sU pulse label, followed by biotinylation and streptavidin bead capture of nascent labelled transcripts. Uniquely, after enrichment of the labelled RNA (without the fragmentation step used in TT-Seq), the 3’ end nucleotide in the nascent transcripts is marked with an artificial poly(A) tail, using bacterial poly(A) polymerase (bPAP). The aim is to allow precise nucleotide resolution mapping of the last base incorporated into the 4sU-labelled nascent transcript. The addition of the artificial poly(A) tail offers a cost-effective and straightforward method for library preparation with standard oligo(dT) primer-based RNA-Seq library kits and commercially available 3’ RNA sequencing kits. Controls are focused on ruling out potential mapping biases and determining the precise source of reads. These include removing reads resulting from priming at internal poly(A) tracks within transcripts using an established computational algorithm (24), an assessment of the proportion of reads from unlabelled transcripts or rRNA, and an assessment of the contribution of polyadenylation by host poly(A) polymerase (hPAP) during the labelling window, similar to those used in a study in *Saccharomyces cerevisiae* (17). Here, 4sU labelling is coupled to direct sequencing of artificially (bacterial poly(A) polymerase-dependent bPAP) polyadenylated RNA 3’ ends, reporting the single nucleotide next to the artificial poly(A) tail as the readout, marking the precise distance RNA polymerases have travelled along the transcription unit and the frequency of pausing at that site by increased read counts, revealing 5’ promoter proximal pauses and the positions of one or more polyadenylation sites (PAS) for RNA polymerase II (RNAPII or PolII) transcribed genes. Controls lacking the bPAP treatment enable detection of host PAP-dependent events during the labelling window, for example, close to the promoter (25), at PROMPTS/PDATs (26) and at nascent PASs. Exact positioning and dependence of the readout on the bPAP poly(A) tail status allow interrogation of transcription rates, RNA synthesis rates, and RNA half-lives at high resolution from single RNA samples. The SNU-Seq protocol for mammalian cells is innovative and straightforward, providing single-nucleotide resolution, good library quality and reproducibility. SNU-Seq is applied to map nascent transcription around genes and enhancers in HEK293 and Hep3B cells, providing new insights into the relationship between chromatin and transcription, including the high number of regions primed for transcription and showing IFNγ-dependent onset of divergent nascent transcription at enhancers and gene promoters. The SNU-Seq readout was complemented with data from sf4sU-Seq, here focused on the nucleotides at the 3’ and 5’ ends of short fragments mapping to the promoter proximal regions, revealing the precise position of the PPP and defining unannotated TSS, particularly for promoter divergent antisense transcripts.

## RESULTS

### Development of single-nucleotide 4sU-Seq (SNU-Seq)

The aim of this work was to use the effectiveness of a 4sU pulse-label to produce cost-effective nucleotide resolution nascent transcriptomes in adherent human cells (Hep3B and HEK293) together with appropriate controls to enable the precise source of each read to be determined (**Fig 1A&S1A**). Optimal labelling times and reproducibility were confirmed by performing TT-Seq in HEK293, HeLa and Hep3B cells (**Fig S1B**). Labelling time was generally a function of growth rate with HeLa cells requiring an 8 min pulse label compared to 10 mins for HEK293 and 10-12 mins for Hep3B cells, sufficient for cells to process the 4sU metabolic label into a triphosphate for incorporation into the nascent transcripts, but short enough to capture primarily nascent, as opposed to processed transcripts, including abundant non-coding transcripts upstream of *KAZALD1* (**Fig S1C**).

**Figure 1.**
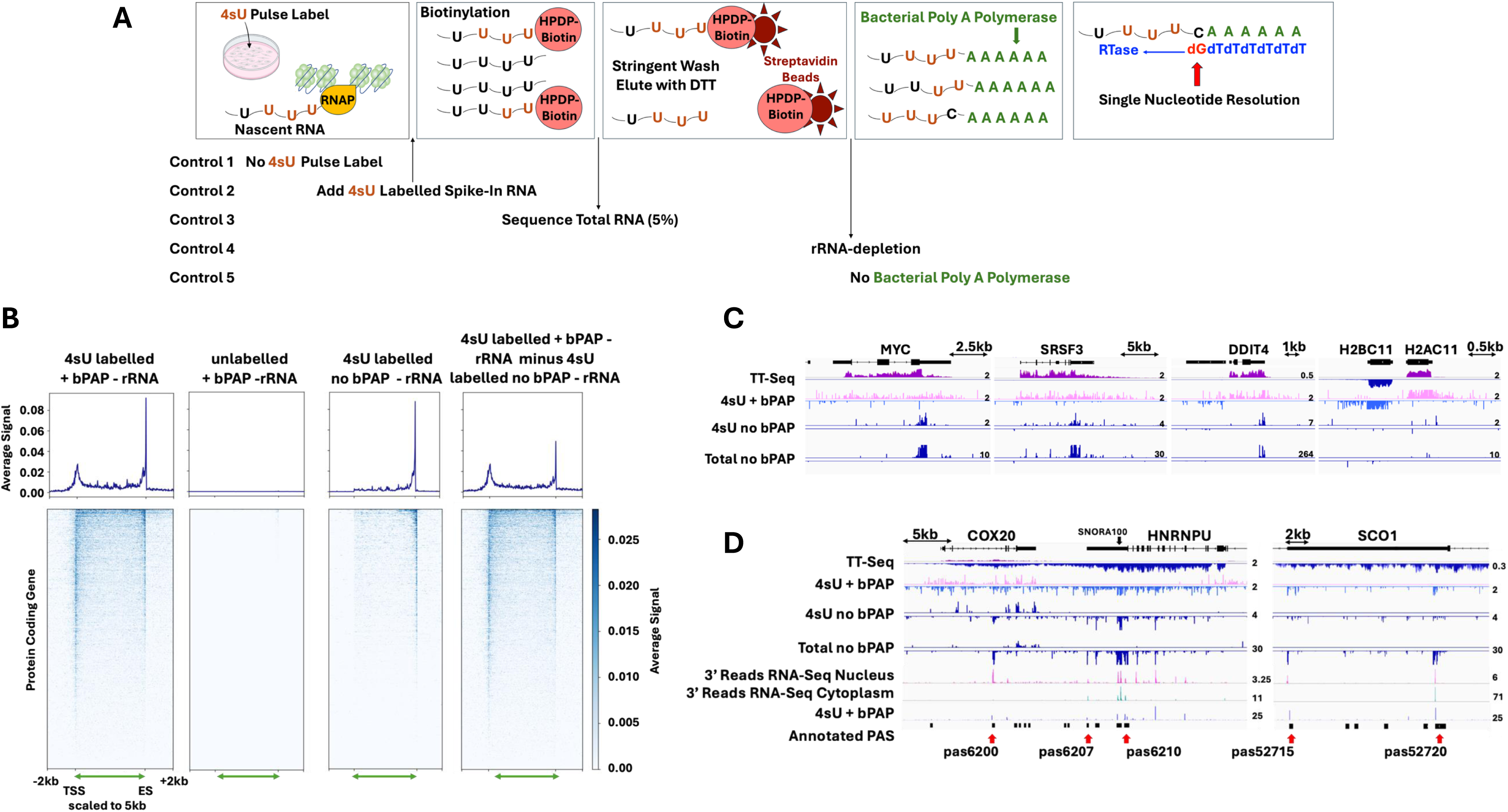
Single-nucleotide resolution 4sU-Seq (SNU-Seq). **A** Schematic flow diagram for SNU-Seq. **B** Metagene profiles and heat maps for 3,883 GENCODE protein coding genes after blacklisting with above threshold signal between the TSS and ES with the transcribed region scaled to 5 kb (green double headed arrow) and the flanking 2 kb up and downstream shown as log_2_ average normalised reads for 4sU labelled or unlabelled samples after biotinylation and selection treated with or without bPAP. Samples were processed as indicated. **C** SNU-Seq detects nascent polyadenylation sites. Snapshots of IGV outputs at the loci indicated. The SNU-Seq output with (4sU + bPAP) or without (4sU no bPAP) bPAP treatment is compared to the total RNA profile (no bPAP), the 3’ RNA-Seq output using 3’ reads (28) and the TT-Seq profile (HEK293, this study). Scale is indicated. **D** Mapping polyadenylation sites (PAS) at *HNRNPU* and *SCO1* in HEK293, with the RNA isolated as three repeats from the nucleus (red) or cytoplasm (blue) (63), and SNU-Seq output (4sU + bPAP, black). Annotated PAS (28) (blue boxes) on the negative strand are indicated with arrows. Scales are indicated for comparison. See also **Figures S1** and **S2**.

In SNU-Seq, a poly(A) tail is added to 3’ end of the nascent transcripts using bacterial poly(A) polymerase (bPAP) and the library sequenced using oligo(dT) priming, comparable to 3’ end RNA-Seq (27,28), reporting just the first base next to the newly added poly(A) tail (**Fig 1A**). Although in both SNU-Seq and TT-Seq nascent transcripts are pulse labelled with 4sU and captured after conjugating HPDP-Biotin and binding to streptavidin beads, they differ as the RNA is not subject to fragmentation in SNU-Seq, as it is in TT-Seq (compared in **Fig S1D**), thus capturing the 3’ end of intact 4sU labelled transcripts. For HEK293 cells, approximately 65% of reads uniquely align, and biological repeats correlate well (**Fig S1E**). Similar read coverage is observed in both HEK293 and Hep3B cells (**Fig S1F**), with no major batch effects observed (**Fig S1G**). To determine the precise source of reads in the SNU-Seq output from HEK293 cells, metagene profiles and heatmaps were prepared for 2,394 genes, including controls outlined in **Fig.1A** after data processing following the scheme in **Fig S1A**.

In the normalised scaled SNU-Seq output, reads can arise from priming by oligo(dT) from internal (A)-rich sequences. As described previously (24), 2.8% of the hg38 genome was masked for genomic (A) residues using the “bedtools intersect -v” algorithm during data processing (**Fig S1A**). After 10 min 4sU labelling, 5% of the samples were processed in parallel as total RNA controls, before or after rRNA depletion (**Fig 1A**). For these samples, metagenes and heatmaps revealed signal predominantly at the 3’ end, expected from oligo(dT) priming from the poly(A) tail of mature mRNA (**Fig.S2A**). Depletion of rRNA improves the resolution of the remaining pre-mRNA signal (**Fig S2A**). When the total RNA samples (Total no bPAP) are treated with bacterial poly(A) polymerase (Total + bPAP), additional signals over the transcribed region are evident, likely representing 3’ ends generated by the synthesis and degradation of transcripts during the labelling window (**Fig S2A**). Subtraction of total no bPAP from total + bPAP reveals the action of bPAP on the transcripts (**Fig S2A**).

The remaining 95% of each sample was biotinylated, the labelled RNA enriched, rRNA depleted, treated with bPAP (4sU labelled + bPAP -rRNA) and compared to the signal obtained when no 4sU is added to cells (unlabelled + bPAP - rRNA) (**Fig 1B**). This indicated virtually no background signal when nascent transcripts were not labelled, compared to the SNU-Seq profile which has reads from the transcription start site (TSS) to the end site (ES) and beyond.

The precise source of the reads in the SNU-Seq output was then examined. Theoretically, reads could also arise from the host cell poly(A) polymerase (hPAP) adenylating nascent 4sU labelled transcripts during the labelling window at polyadenylation sites (PAS), as observed in a similar technique developed for *Saccharomyces cerevisiae* (17). To control for this, the SNU-Seq libraries prepared after treatment with bacterial poly(A) polymerase (4sU labelled + bPAP -rRNA) were compared to the profiles for 4sU-labelled transcripts without bPAP treatment (4sU labelled no bPAP - rRNA) (**Fig1B**). In both samples, the predominant signal at the 3’ region maps to the transcript end site (ES), consistent with poly(A) tails added during the labelling window by the host poly(A) polymerase (hPAP).

Subtraction of the signal in the 4sU labelled no bPAP samples (host PAP only) from the SNU-Seq (4sU labelled + bPAP) samples (both host and bacterial PAP) reveals the bPAP- dependent signal in SNU-Seq. This leaves peaks primarily in the promoter proximal region and at the PAS (**Fig 1B&S2B**). The SNU-Seq output (4sU labelled + bPAP) was used for all subsequent analysis, as this includes additional potentially valuable information on polyadenylation site usage. No major batch effects are observed when comparing seven 4sU-labelled samples treated either with or without bPAP (**Fig S2C**).

Screenshots of IGV SNU-Seq profiles with controls at six genes were used to illustrate two features evident at 3’ regions (**Fig 1C,D**). First, increased read density and spikes in the SNU-Seq (4sU + bPAP) overlap with mapped PAS evident in the SNU-Seq control (4sU no bPAP, indicative of the action of host PAP during the labelling window) and in the total RNA (Total no bPAP). Generally, these spikes are not evident in the TT-Seq (HEK293 cells, this study), although, as previously described (15), a drop in read density occurs around the annotated PASs. Second, the SNU-Seq readout (4sU + bPAP) often extends way beyond the main PAS used in the mature mRNA (total RNA). Further analysis at *HNRNPU* and *SCO1* revealed that this may reflect cleavage and polyadenylation occurring at alternative annotated PAS often located far downstream, for example, the spike at pas6200, >10kb downstream of *HNRNPU* and the two PAS (pas52715 and pas52720) at *SCO1* (**Fig 1D**). Details analysis of 3’ reads output in HEK293 cells when RNA is separated into nuclear and cytoplasmic fractions reveals overlap with the 3’ reads output in the nucleus and the spikes in SNU-Seq profile. However, the profile of reads in the cytoplasm is markedly different, often reflecting preferential accumulation of transcripts using the proximal PAS. Finally, no signal in the control (4sU no bPAP; indicative of the action of the host poly(A) polymerase) is evident at histone genes (*H2BC11*, *H2AC11*) whose 3’ ends are not subject to polyadenylation (**Fig 1C**). Thus, the SNU-Seq output also contains information about usage of polyadenylation sites during the labelling window.

As SNU-seq is a simple technique, comparable to 3’ end RNA-Seq (27,28), it potentially offers real advantages compared to other methods for assessing nascent transcription at genes and enhancers. Metagenes, heatmaps and a genomic snapshot in IGV around *CERS6* and *DDIT4* illustrate the differences in the output signals generated by SNU-Seq (This study), TT-Seq (This study and (29)), PRO-Seq (29), and mNET-Seq (30) in HEK293 cells (**Figs 2A,B&S3**). SNU-Seq generally shows a low variance in the signal height and provides a flat profile, with the density of reads reflecting the amount of label incorporated during the labelling window and spikes at the promoter proximal region, around polyadenylation sites and pauses. The SNU-Seq output is distributed over both exons and introns, supporting effective capture of nascent transcription before extensive pre-mRNA processing (**Figs 2B&S3**). There is a notable difference between SNU-Seq and the two TT-Seq profiles, particularly the inability of TT-Seq to resolve the 5’ and 3’ peaks (**Fig 2A,B**). Both PRO-Seq and mNET-Seq resolve the spike at the promoter proximal region but not at the position of the PAS. The SNU-Seq signal correlates well (Spearman correlation 0.672) with mature transcript levels at 12,303 genes (31) in HEK293 cells (**Fig 2C**) and accurately reflects transcription elongation profiles established for the TT-Seq protocol in K562 cells (15,32) (**Fig 2D**). Finally, the ability of the four techniques to distinguish eRNAs characterised enhancers around *DDIT4* in HEK293 cells was examined, using the chromatin signature to map open chromatin (ATAC-Seq) surrounded by H3K27ac (E, red arrows, **Fig S3B**). While divergent transcription is evident in the SNU-Seq read out, eRNA are not reliability detectable in the TT-Seq, PRO-Seq or mNET-Seq outputs. For mNET-Seq this reflects the specificity of antibodies used in the immunoprecipitation step. In PRO-Seq, eRNA at the upstream enhancer are evident but not the two downstream enhancer elements, even when the scales are optimised relative to *DDIT4* signal. Thus, SNU-Seq can generate high-quality data for nascent transcripts in human cells, with multiple features of nascent transcripts in one simple readout. Before validating these features, particularly the promoter proximal pause (PPP) and eRNAs, the SNU-Seq data were used to calculate transcription parameters in HEK293 cells.

**Figure 2.**
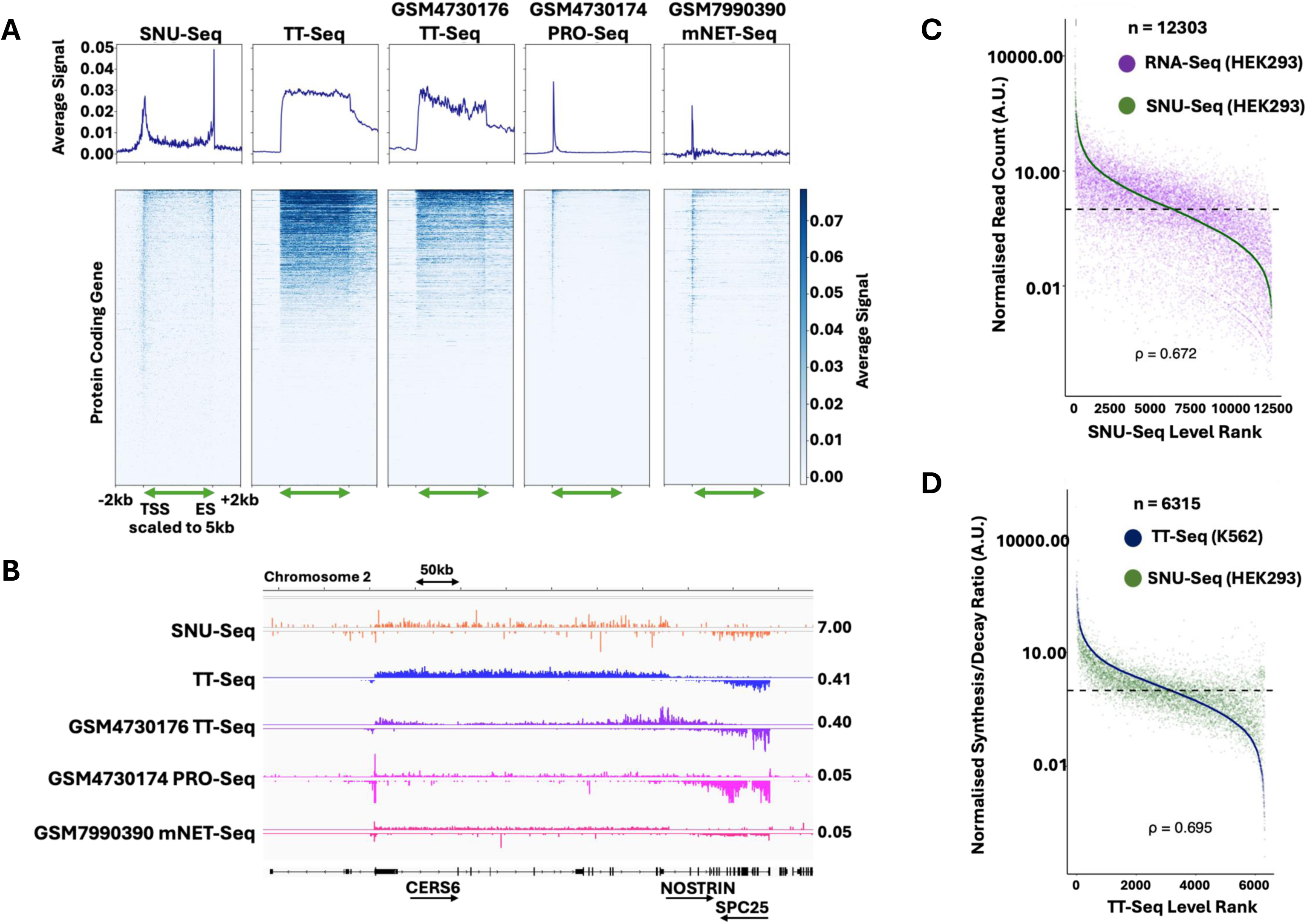
Comparison of SNU-Seq and other methods for assessing nascent transcription. **A** Metagenes and heat maps of SNU-Seq and TT-Seq plotted on the same scale, illustrating the promoter proximal peak and PAS in SNU-Seq and compared to TT- Seq, PRO-Seq and mNET-Seq (sources indicated) also in HEK293 cells. **B** IGV screenshot around *CERS6* displaying a comparison of SNU-Seq (10 min 4sU pulse in HEK293, this study n = 3), TT-Seq (10 min 4sU pulse in HEK293, this study n = 2), TT-Seq, PRO-Seq and mNET-Seq also in HEK293 cells from sources indicated. **C** Correlation between SNU-Seq (this study) and published RNA-Seq in HEK293 cells (31). The Spearman correlation coefficient (rho) is indicated. **D** Ranked correlation plot of the synthesis-to-decay ratios between SNU-Seq (this study) and the original TT-Seq dataset in K562 cells (15). The Spearman correlation coefficient (rho) is indicated. (**See also Figure S3**).

### Synthesis and decay rates

After thresholding and removing reads in ENCODE blacklisted regions, the synthesis rate (SR) and decay rate (DR) were calculated for 2,394 genes in the annotation subset: DR = - (1/time) x log(1-RNA_4sU_/RNA_total_), and SR = RNA_total_ x DR, where time is 10 min, RNA_4sU_ is the normalised SNU-Seq counts with bPAP treatment, exploiting the total RNA signal (no bPAP) and the 4sU labelled (+ bPAP) nascent RNA signal from the same samples (15,32). The density of reads over the gene body directly relates to their synthesis rates (Kendall’s T = 0.99) (**Fig 3A**). Based on an elbow analysis of the calculated synthesis rates, k-means clustering (k = 3) was applied and used to classify genes into fast, middle and slow groups. In each group, fast synthesis rates have higher spike-in normalised counts and lower decay rates, like parameters determined in *S. cerevisiae* (33) (**Fig 3B-D**).

**Figure 3.**
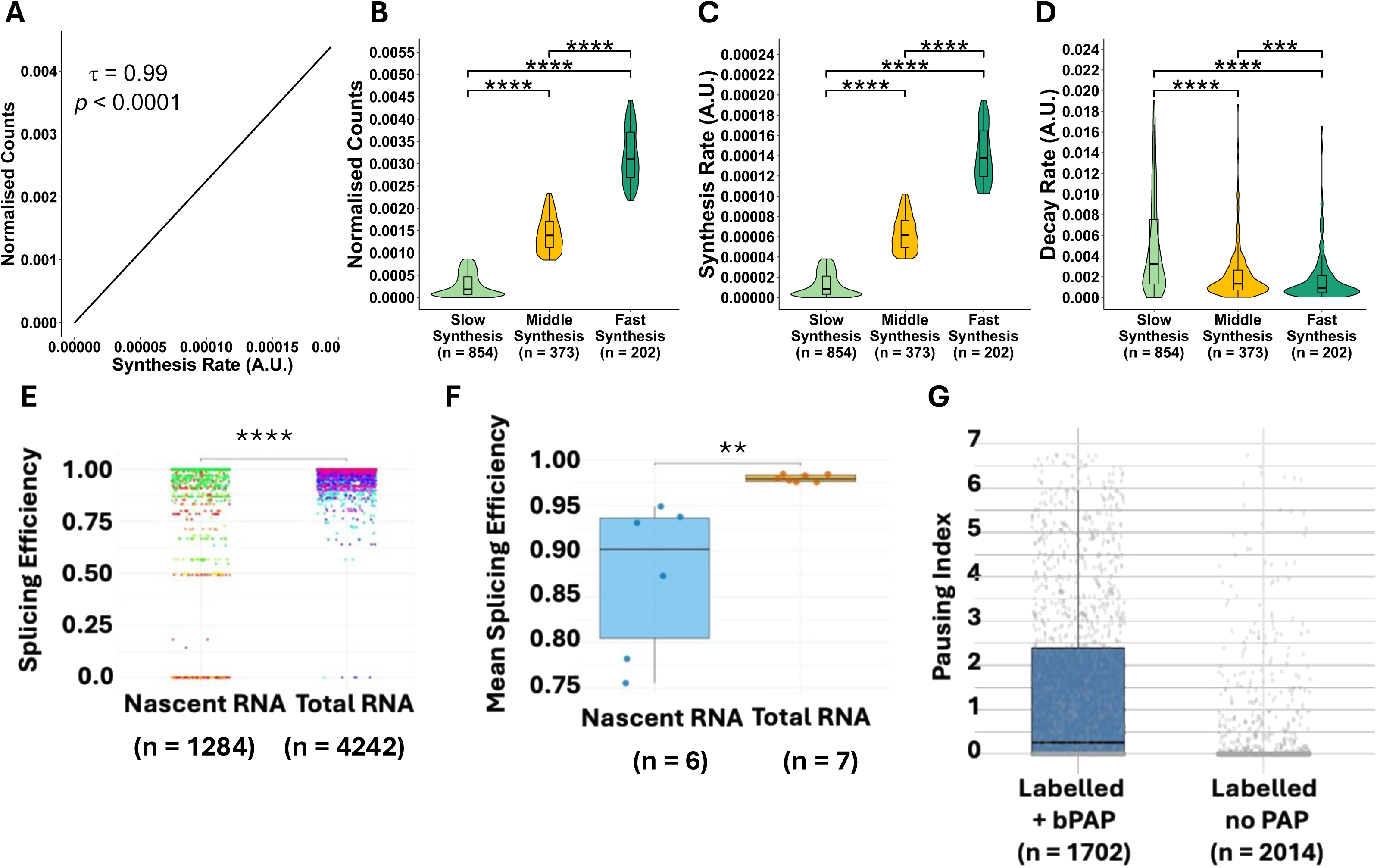
Transcription parameters derived from SNU-Seq data. **A** Correlation between synthesis rate and spike-in normalised counts, with Kendall’s tau correlation coefficient (τ = 0.99) and p-value (p < 0.0001). **B-D** Distributions of synthesis rate, decay rate, and spike-in normalised counts grouped by synthesis rate clusters (k = 3). The 1,027 active genes are distributed across three transcriptional activity clusters: Slow Synthesis (n = 408), Middle Synthesis (n = 403), and Fast Synthesis (n = 216). The significance of differences between clusters was assessed using the Wilcoxon test, with **** denoting p < 0.0001. **E**-**F** Splicing efficiency and mean splicing efficiency derived for total RNA preparation (total RNA; n = 7) compared to 4sU-labelled and selected nascent RNA (nascent RNA n = 6) in HEK293 cells. **** p<0.0001, ** p < 0.01. **G** Pausing index comparing 4sU labelled RNA treated with bPAP or without bPAP.

### Splicing efficiency in nascent transcript output

To assess the level of pre-mRNA processing captured during the labelling window by SNU- Seq, the primary 3’ end-Seq output obtained from the total RNA libraries (n = 7) and the SNU-Seq (nascent RNA) libraries (n = 6) were used to calculate the splicing efficiency from the ratio of spliced to unspliced reads at splice junctions. Using the SPLICE-q tool (34), the mean splicing efficiency (SE) score was computed by averaging all measured splicing events in the total RNA libraries and SNU-Seq libraries prepared from the same samples (**Fig 3E, F**). The results indicated that SNU-Seq captures more transcripts with introns (nascent RNA) in comparison to reads from the total RNA, which also includes fully processed transcripts, as expected.

### 5’ end Pausing Index

Metagene profiles and IGV snapshots indicate read peaks near the TSS of transcription units, likely representing promoter proximal pausing of initiated RNA polymerase (see **Fig 1B**). This pause is generally only captured in 4sU labelled samples when treated with bPAP, as the output from 4sU-labelled samples not treated with bPAP primarily captures reads due to polyadenylation of pre-mRNAs undergoing 3’ end formation during the labelling window. The read counts within the TSS proximal window (-50 bp to +200 bp) were compared to those from +200 bp to the start of the 3’ UTR for the two data sets and indicate a much higher ratio in the bPAP-treated nascent transcripts (**Fig 3G**). Next, the sensitivity of 4sU labelling was exploited to isolate and characterise the short transcripts whose 3’ ends are found within the first 200 nt of genes.

### Using size fractionated 4sU-Seq (sf4sU-Seq) to explore the TSS-proximal peak in the HEK293 nascent transcriptome

Heatmaps suggest that many genes have a read distribution consistent with short 4sU labelled nascent transcripts in the region proximal to the promoter (see **Fig 1B**). Generally, these short transcripts are not subject to polyadenylation by host PAP as there are no significant reads in the control samples (4sU labelled no bPAP) (see **Fig 1B**). To investigate these short transcripts, the biotinylated, 4sU-labelled RNA was subject to electrophoresis on a polyacrylamide gel, similar to the approach used for size-fractionated NET-Seq in *S. cerevisiae* (5) (**Fig 4A**). When compared to the total RNA from HEK293 cells, the nascent thio-labelled RNA reproducibility yields a clear band at approximately 50-100 nt and a smear from ≍ 600 nt upwards on gels indicating that the nascent 4sU-labelled transcripts are not subject to extensive degradation, which was confirmed once the short fraction was sequenced (**Fig S4A, B**). The short RNA was isolated (**Fig S4C, D**) and subject to linker ligation at both the 3’ end (75 nt) and 5’ end (54 nt) after decapping with RNA 5’ pyrophosphohydrolase (**Fig S4E**) and sequenced from both ends. The coverage showed more the 86% of the library aligned to the human genome (**Fig S4F**), and after processing, allows the first 5’ and the last 3’ nucleotides of the gel-purified nascent transcripts to be reported, yielding base pair resolution.

**Figure 4.**
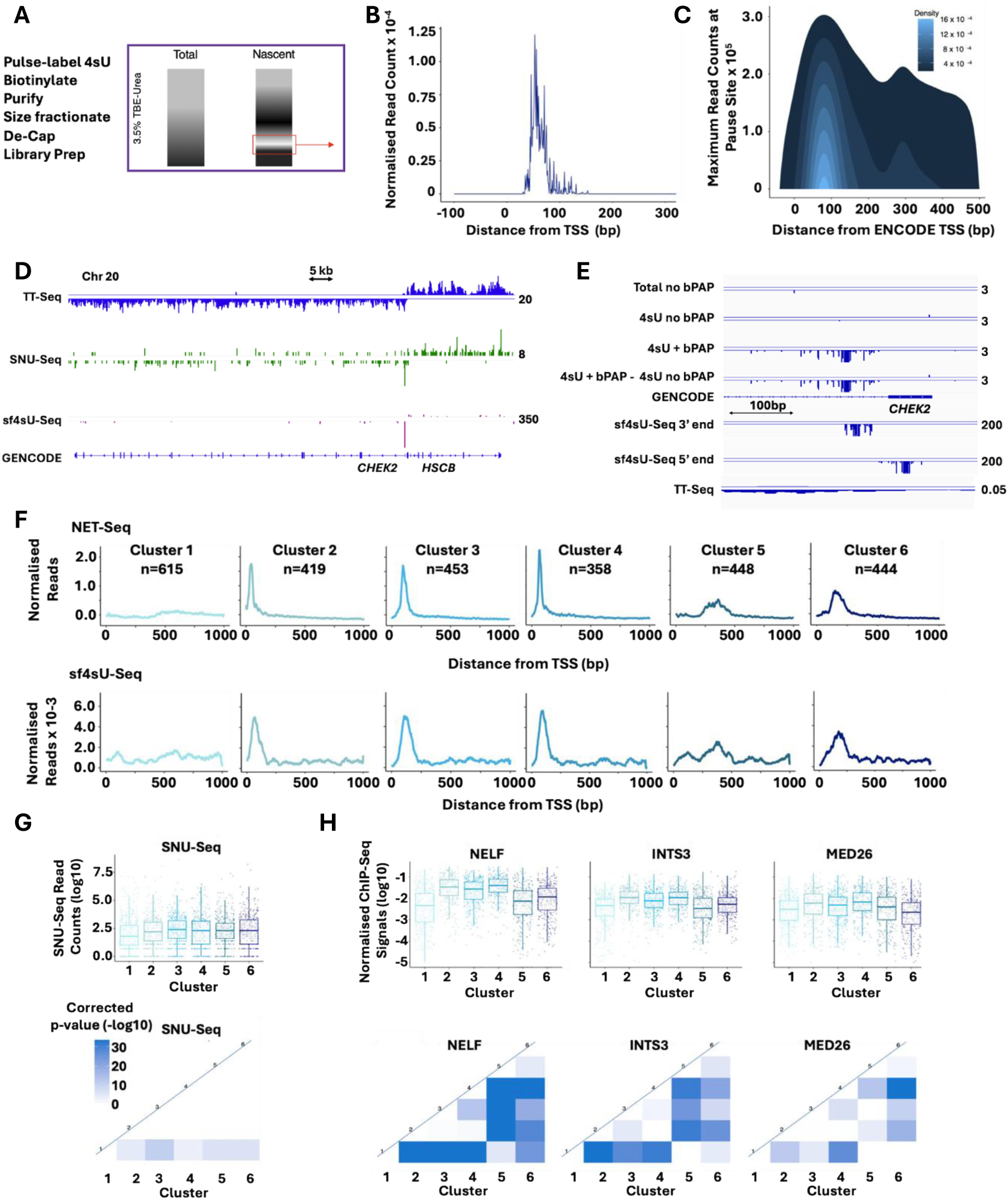
Size fractionated 4sU-Seq (sf4sU-Seq) captures the promoter-proximal pause. **A** Workflow for 4sU-Seq with size-fractionation including thio-labelling, biotinylation, and streptavidin purification, followed by a gel-based size fractionated step. After purifying RNAs of a size range between 50 and 100 nucleotides and decapping, adapters were added immediately for library preparation, retaining single-nucleotide resolution at both the 5’ and the 3’ ends. **B**. Normalised metagene of size-fractionated 4sU-Seq reads around human TSSs (n = 9,336). **C** 2-dimensional density plot showing the distribution of the maximum read number of each gene against its location relative to the ENCODE transcription start site. Density refers to the number of occurrences per pixel (n = 19,874). **D** IGV screenshot displaying TT-Seq (HEK293, this study), SNU-Seq (HEK293, this study), and size- fractionated 4sU-Seq (HEK293, this study) profile around the *CHEK2* locus on chromosome 20. Scales are chosen to highlight read spikes. **E** Zoomed in view at the 5’ region of *CHEK2* with additional SNU-Seq controls demonstrating no polyadenylation at the promoter proximal signal, as indicated, and output from sf4sU-Seq at the 5’ and 3’ ends of the isolated fragments. Scales are chosen to highlight clusters. **F** Generation of k-means clusters (k = 1 to k = 6) using human (HeLa) mNET-Seq (2). The same cluster indices for each gene were applied to sort size-fractionated 4sU-Seq data (HEK293 cells). Metagenes are normalised to make every gene contribute equally. **G-H** Statistical analysis of SNU-Seq, NELF, integrator, and mediator occupancy within gene clusters. Boxplot diagrams for NELF, INTS3, MED26, and SNU-Seq levels at the 5’ end of genes for each cluster (300 bp window for ChIP-Seq and 1,000 bp window for SNU-Seq). ChIP-Seq data were normalised to make each gene contribute equally. For SNU-Seq, the raw counts were used. All data were log-transformed. Triangular heatmaps of between-cluster significance for NELF, INTS3, MED26, and SNU- Seq levels. p-values were calculated using the Wilcoxon-rank sum test and corrected using the Bonferroni method. (See also **Figures S4 and S5**).

Calibrated metagenes and density maps were created to examine the genome-wide distribution of reads in the sf4sU-Seq libraries (**Fig 4B, C**). The highest density of 3’ end read signal is located around 60-80 bp downstream of the TSS (centred around 63 nts), and a minor density peak is around 300 bp downstream of the TSS. These two peaks may represent the promoter-proximal and promoter-distal pausing sites. Alternatively, they may represent major unannotated transcription start sites (uTSSs) downstream of the annotated site.

At individual genes, for example *CHEK2*, the 5’ TSS proximal peak is evident in SNU-Seq and in the sf4sU-Seq, but not in the TT-Seq data (**Fig 4D**). The 5’ end of the sf4sU-Seq fragment is consistent with the TSS mapped using TT-Seq and the 3’ ends overlap with the signal in the SNU-Seq readout at the pause site (**Fig 4E**). The SNU-Seq control (4sU no bPAP) confirms that the promoter proximal short transcripts are generally not subject to polyadenylation (**Fig 4E**), although there are exceptions, for example *ZNRF3*, where both divergent short transcripts, including the PDAT/PROMPT (26), have a signal in the control (4sU no bPAP) suggesting polyadenylation by hPAP (**Fig S4G**).

The TSS proximal signal from the sf4sU-Seq data (3’ ends) was compared to mNET-Seq data, which captures all forms of RNAPII, including the promoter proximal pause (2), using an unsupervised machine learning approach. First, k-means clustering (k = 6) was performed on the shape of human NET-Seq profiles for 2,748 genes to identify clusters of genes with different shapes of promoter proximal profiles (**Fig 4&S4H**). This yielded one cluster with a flat metagene (cluster 1), three clusters with a sharp peak directly downstream of the TSS (clusters 2, 3, and 4), one cluster with a slightly broader peak (cluster 6), and one cluster with a broad peak much further into the gene body (cluster 5). There was no difference in the levels of transcription contributing to the shape of clusters, as SNU-Seq reads over the first 1,000 nucleotides are similar in all six clusters (**Fig 4G**). In addition, there is no evidence for a DRB-sensitive promoter proximal pause in the flat class, although the “peaky” signal in the remaining clusters (k = 4; NET-Seq) increases in size over time of DRB-treatment as release from the pause is inhibited (**Fig S4I**). sf4sU-Seq data were plotted for the genes in each cluster, and remarkably, the patterns match those in the NET-Seq data, suggesting that the position of the promoter proximal pause is captured by reads at the 3’ end of the short fragments isolated in sf4sU-Seq (**Fig 4F**).

To provide additional evidence for the nature of the promoter proximal signal, ChIP-Seq data for NELF, a pausing factor (35), INTS3, a subunit of integrator (36), linked to the pausing signal by premature termination, and MED26 (37,38) with a role in recruitment of the super elongation complex (SEC) to polyadenylated genes (39–43) was used to validate these peaks as pauses or sites of early termination (**Fig S4H**). For all three factors, the profiles reflect the NET-Seq and sf4sU-Seq profiles, although only the mediator ChIP-Seq profile (MED26) shows the dual peaks in cluster 5, possibly representing alternative start sites. This was confirmed using CoPRO, a nuclear-run on variation of PRO-Seq that enriches for 5’ end states of nascent RNAs (44) and ATAC-Seq data from HEK293 cells (45) which revealed that the peak in cluster five is shifted significantly further downstream compared to the other clusters (adjusted p-value < 3.5 x 10**^-^**^5^) (**Fig S4H**). The distribution of the levels of NELF, INTS3, and MED26 over the first 300 nt was assessed (**Fig 4H**). NELF shows the greatest enrichment in peaky clusters 2, 3 and 4, compared to INTS3 and MED26. This suggests that levels of NELF are the best predictor of a promoter proximal pause, and that the majority of the 3’ end reads in the sf4sU-Seq data mark the position of the PPP.

A simple machine learning algorithm (logistic regression) was used to assess whether NELF, integrator, or mediator levels alone are predictive of which cluster a gene may belong to: pausy; clusters 2-4 or non-pausy; cluster 1 (k=4) (**Fig S5A-F**). NELF shows the strongest predictive power, compared to integrator or mediator subunits, in terms of being able to classify mNET-Seq-based clusters as pausy or non-pausy. This supports the signals in the SNU-Seq data as primarily reflecting pausing around 60-80 nt downstream of the TSS. As INTS3 also shows enrichment, some of the 3’ end signals in the sf4sU-Seq could arise due to integrator-dependent early termination of transcription (25,46). To characterise the regulatory context, the calibrated pausing signal was obtained by dividing the summed sf4sU-Seq signals for each gene (50 – 150 bp downstream of the TSS) by the sum of TT-Seq reads for each gene (0 – 500 bp downstream of the TSS, **Fig S5G**). Gene ontology analysis of the top and bottom deciles of genes ranked by their relative pausing signal revealed that genes exhibiting pausing are involved in primary metabolic processes, cell-cycle regulation and DNA-damage checkpoints (**Fig S5H**). This agrees with previous analyses on promoter-proximal pausing based on Pol II ChIP-Seq (47). Because of the single-nucleotide resolution, as well as the high sensitivity of this approach, highly statistically significant results on the gene ontology of promoter-proximal pause were generated compared to the ChIP-Seq approach, which generates low-resolution Pol II positioning profiles rather than precise information about the act of transcription. Thus, sf4sU-Seq validates the 5’ peak in the SNU-Seq metagenes as the promoter proximal pause (PPP) located 60-80 nt from the TSS.

### Using size fractionated 4sU-Seq (sf4sU-Seq) to discover unannotated TSS in the HEK293 transcriptome

Based on the 5’ end signal at the short fragments in the sf4sU-Seq data, a pipeline was developed to facilitate annotation of transcription start sites and to discover new unannotated TSS (**Fig 5**), using the same principle as for START-Seq annotations (48–50). First, only 5’-end signals that overlap with ATAC-Seq peaks in HEK293 cells were used (45) (**Fig 5A,B**). Second, the Paraclu TSS-clustering algorithm was used to group closely positioned TSS, as used in CAGE-Seq (51–53). The sf4sU-Seq based clusters are predominantly shorter than 10 bp (**Fig 5C**) and the cluster-centres map to ENCODE TSS annotations as expected (**Fig 5D**). TSS candidates for which the TT-Seq signal (in HEK293 cells) in the first 1,000 bp downstream of the TSS candidate was lower than 5 times the signal of the 1,000 bp preceding the TSS candidate were filtered out (**Fig 5E**). This resulted in 4,383 candidate TSSs distributed widely over the genome (**Fig 5F**) and associated with active transcription units mapped using TT-Seq (**Fig 5G**). Finally, TSSs were assigned as previously annotated (observed TSSs or obsTSSs) or unknown/novel/unannotated TSSs (uTSSs) using the open-source code TSScall (50) (**Fig 5H**). The vast majority of the 2,955 active TSSs map around ENCODE annotated TSSs (**Fig 5I**). Many of the 1,428 newly-identified TSSs are predominantly divergent (antisense TSS, associated with PDATs or PROMPTS (54)) and are located between 100 and 500 bp upstream of the sense TSSs (**Fig 5J**), and shown in metagenes aligned to ENCODE TSS (**Fig 5K**).

**Figure 5.**
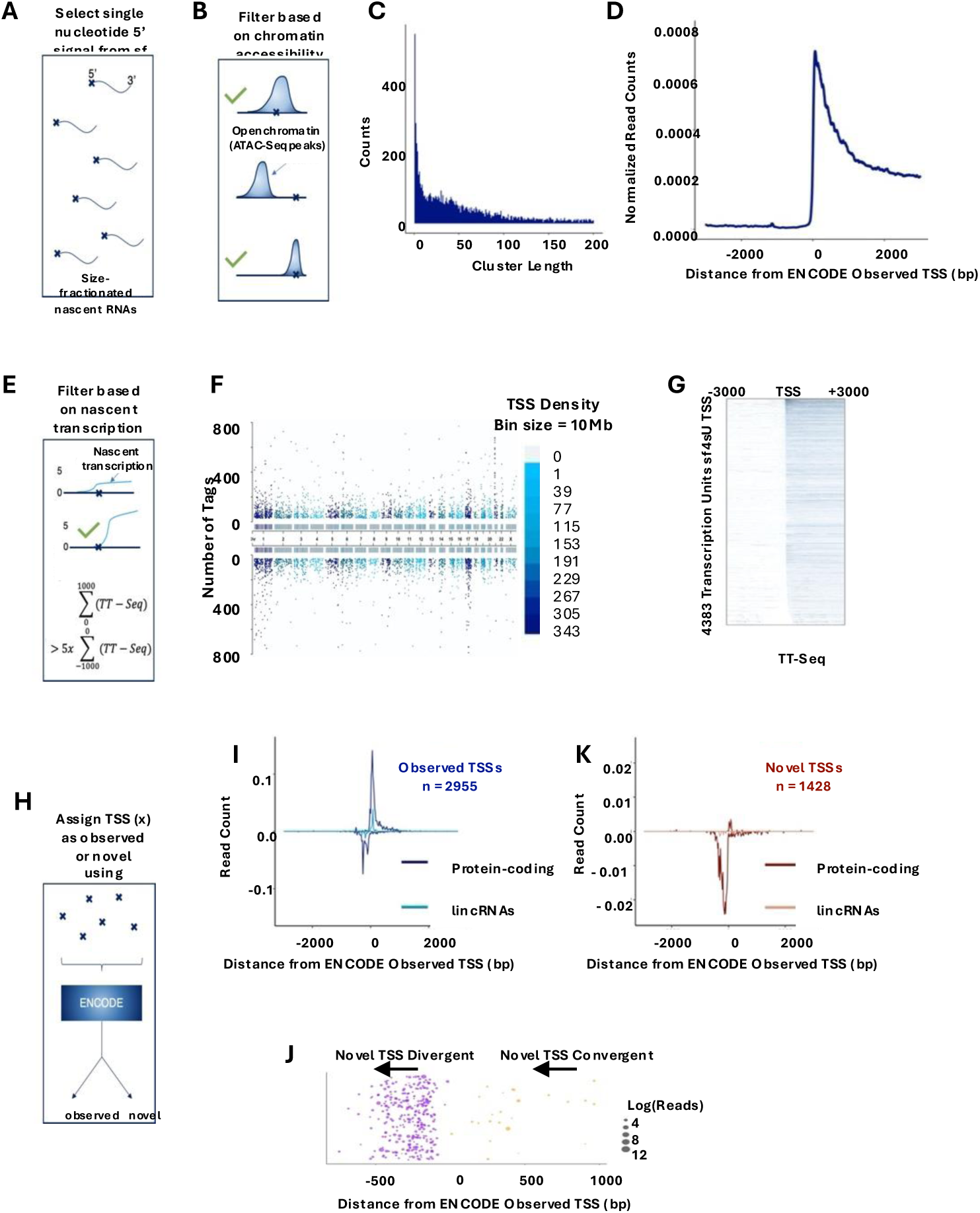
**sf4sU-Seq-based TSS characterisation**. **A, B, E, H**. A simplified overview of the steps involved in the TSS annotation workflow linked to the data output. 5’ end signals of size- fractionated 4sU-Seq were filtered through ATAC-Seq peaks and, after clustering into TSS- clusters, filtered based on a minimum 5-fold increase in the TT-Seq signal immediately downstream of the TSS candidate. The verified TSSs were then classified as either previously annotated (obsTSSs) or novel / unannotated TSSs (uTSSs). **C** Clustering of sf4sU-based TSS candidates showing a histogram of the cluster lengths. **D** A metagene of the full-length clusters around ENCODE transcription start sites. The signal is normalised so that each locus contributes equally to the metagene by making the sum of reads for each cluster equal to 1. **F** Manhattan plot of sf4sU-Seq-based TSS annotations. The plot shows the location of each verified TSS annotation on the respective strand (+ or -) and chromosome. The y-axis corresponds to the number of reads that were associated with the original 5’ end signal. The bars show the density of TSSs within each chromosome on each strand. **G** A heatmap of TT-Seq around sf4sU-Seq-based TSS annotations (signal range 0 - 500 reads), ordered by number of reads from 1 bp downstream of the TSS in descending order and displayed in a 6 kb window around the TSS. **I** Metagene of already annotated TSSs (obsTSSs, n = 2,955) around protein-coding or lincRNA ENCODE TSSs. **J** Categories of sf4sU-Seq-based TSSs that have not been annotated previously are classified as either convergent or divergent if they are in the vicinity of already annotated TSSs and on the opposite strand. The size of the data points corresponds to the log-transformed read count. **K** Metagene of novel/unannotated TSSs (uTSSs, n = 1,428) around protein-coding or lincRNA ENCODE TSSs.

In conclusion, size fractionated 4sU-Seq is a viable method for transcription start site annotation, here relying on a strict set of filters, such as ATAC-Seq peaks, and a 5-fold increase in TT-Seq signal, which leads to a relatively small set of TSSs. Because of these filters, these TSSs belong to transcription units that are transcriptionally active and found in open and accessible chromatin. 4sU-based approaches, such as SNU-Seq, seem particularly suitable when interested in detecting non-coding and antisense transcription. This holds true for 4sU-based TSS annotations obtained through size fractionated 4sU-Seq, too, given the 1,428 unknown TSSs that are antisense to the TSSs of already annotated protein coding genes. To explore and expand these findings, we used SNU-Seq combined with chromatin analysis (ATAC-Seq, H3K27ac, H3K4me3, CTCF) to assess transcription from promoters and enhancers in Hep3B hepatocarcinoma cells, which, like hepatocytes, are well characterised (55).

### Using SNU-seq and chromatin analysis to characterise functional elements in the Hep3B genome

To demonstrate that SNU-Seq (4sU labelled + bPAP; nascent RNA) can detect nascent transcripts at enhancers (eRNAs) in Hep3B cells, previously characterised functional enhancers interacting with the enhancer-looping factor LDB1 in hepatocytes (56) were visualised in IGV (57). In addition to SNU-Seq, mature mRNA samples were prepared, tagmentation was used to map regions of open chromatin (ATAC-Seq, n = 3), and ChIPmentation was used to monitor regions enriched for H3K27ac (n = 2), H3K4me3 (n = 2) and CTCF (n = 3).

At *SLC2A2*, four hepatocyte enhancers E1-E4 (56) are coincident with open chromatin, peaks of H3K27ac and divergent nascent transcription in Hep3B cells, which within the transcribed region are easiest to identify on the antisense strand (**Fig 6A**). Similarly, at *ALCAM,* four potential enhancers are detectable in Hep3B cells, including 3 previously characterised in hepatocytes within the transcribed region (56) (**Fig S6A**). At 3 regions around *HNF4A,* encoding a liver-specific transcription factor (**Fig S6B**), at 4 regions around *DDIT4* (**Fig S6C,** see also **Fig S3B**) and at *STAT1* (**Fig 6B**), regions with similar characteristics are evident. To confirm that these regions are functional enhancers, Capture-C anchored to the *STAT1* promoter (58) was used to identify a long-range interaction to the 5.5URR upstream enhancer (**Fig 6C-E**). Thus, open chromatin (ATAC-Seq), peaks of H3K27ac and divergent nascent transcription extending > 1 kb in each direction are evident at the *SLC2A2*, *STAT1*, *ALCAM*, *DDIT4* and *HNF4A* enhancers and promoters.

**Figure 6.**
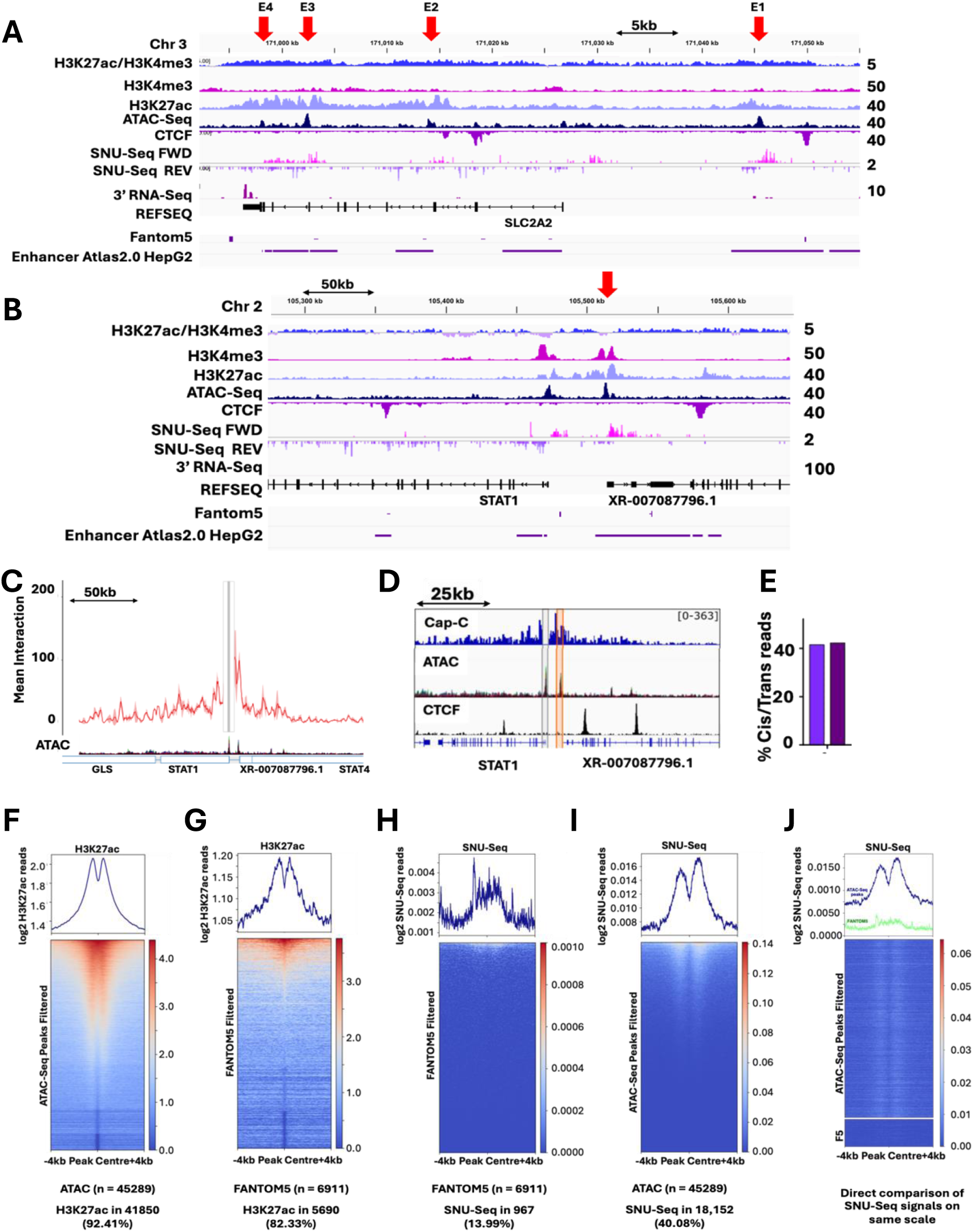
The chromatin environment and nascent transcription in Hep3B cells. A,B. Snapshots in IGV showing chromatin features and nascent transcription around two loci selected as containing previously characterised enhancers (*SLC2A2* **A**; *STAT1* **B**) and known to be expressed in Hep3B cells. Known enhancers are indicated by red arrows. **C-E** Normalised Capture-C data showing mean reads for the *STAT1* promoter (n = 2). Reads were normalised based on the number of cis reads per 100,000 reads. The data is smoothed based on mean reads within a 2 kb window. ATAC-Seq data is shown below the Capture-C data (**C**). **D** Significant differential Capture-C interactions between the *STAT1* promoter and enhancer. Reads are counted per NlaIII-digested fragment. The top tracks show cis-normalised mean data. Normalised ATAC and CTCF tracks are shown. The position of the *STAT1* probe is highlighted in grey, and the putative *STAT1* enhancer element is highlighted in orange. Bar chart showing the percentage of cis reads over the total reads for each Capture-C sample and repeat. Similar ratios indicate similar library qualities. **F-J** Metagenes and heatmaps showing H3K27ac distribution (**F, G**) or SNU-Seq signals (**H, I**) around all ATAC-Seq peaks (**F, I**) or at FANTOM5 annotations (**G, H**) centred around the peaks and extending 4 kb up or downstream. The proportion of total features enriched with H3K27ac or SNU-Seq reads over zero is shown below. **J** A direct comparison of the SNU-Seq reads at all ATAC-Seq peaks and FANTOM5 enhancers plotted on the same scale. Data in **J** were sorted by the ratio of H3K27ac/H3K4me3. **See also Figure S6.**

To examine how frequently these features occur, a genome-wide analysis was conducted by identifying regions of open chromatin (derived experimentally using ATAC-Seq) (n = 64,536), enhancers (FANTOM5, n = 63,285) or promoters (GENCODE annotated TSS -1 kb to +200 bp, n = 70,611). As many of these features overlap with each other, the datasets were filtered by retaining only those on autosomal chromosomes (chr1-22) and identifying any similar features closer than 5 kb and removing both. For ATAC-Seq annotations located up to 1 kb from a GENCODE gene, only those 5 kb apart from any other ATAC-Seq peaks were kept, using this gene’s strand information. This yielded 17,367 annotations on the forward (FWD) strand and 16,570 on the reverse (REV) strand. If features were > 1 kb away from a GENCODE gene, only those at least 5 kb from other peaks were kept and annotated as intergenic features (n = 13,434). All features, regardless of position, and at least 5 kb apart from each other, constituted the third group, n = 45,289 (all). FANTOM5 enhancers were filtered, retaining those at least 5 kb from any GENCODE annotated gene and 5 kb apart from each other, yielding 6,911 features for analysis (**Table 1**). The filtered data were stratified into subsets based on overlaps, and the number of overlapping features was determined (**Table 1**).

**Table 1.** Subsets of ATAC-Seq peaks, associated histone modifications and the SNU- Seq output in Hep3B cells. Relates to. **Figure 7**.

| Subset | Signal | Total number of regions | Number of regions with signal | Percent of regions with signal |
| --- | --- | --- | --- | --- |
| ATAC_peaks_filtered | K27ac_concat | 45289 | 41850 | 92.41 |
| ATAC_peaks_filtered | K4me3_concat | 45289 | 41867 | 92.44 |
| ATAC_peaks_filtered | SNUseq_concat | 45289 | 18152 | 40.08 |
| ATAC_peaks_filtered_fwd | K27ac_concat | 17367 | 16208 | 93.33 |
| ATAC_peaks_filtered_fwd | K4me3_concat | 17367 | 16154 | 93.02 |
| ATAC_peaks_filtered_fwd | SNUseq_concat | 17367 | 8430 | 48.54 |
| ATAC_peaks_filtered_rev | K27ac_concat | 16570 | 15474 | 93.39 |
| ATAC_peaks_filtered_rev | K4me3_concat | 16570 | 15371 | 92.76 |
| ATAC_peaks_filtered_rev | SNUseq_concat | 16570 | 8279 | 49.96 |
| ATAC_peaks_filtered_unstranded | K27ac_concat | 13434 | 12166 | 90.56 |
| ATAC_peaks_filtered_unstranded | K4me3_concat | 13434 | 12369 | 92.07 |
| ATAC_peaks_filtered_unstranded | SNUseq_concat | 13434 | 2927 | 21.79 |
| FANTOM5_filtered | K27ac_concat | 6911 | 5690 | 82.33 |
| FANTOM5_filtered | K4me3_concat | 6911 | 5864 | 84.85 |

Focused on the peak of open chromatin signals from the genome-wide ATAC-Seq data (all), heatmaps and metagenes of the concatenated and log_2_-transformed values for SNU-Seq, H3K27ac and H3K4me3 were plotted (**Fig 6F-J&S6D-H**). 80-90% of accessible sites in chromatin are subject to detectable levels of H3K27ac (**Fig 6F**) or H3K4me3 (**Fig S6D**) including regions annotated as FANTOM5 enhancers with open chromatin (**Fig 6G&S6E**). Metagenes and heatmaps were used to display the SNU-Seq signal at each of the genomic regions and reveal the lowest number of reads at FANTOM5 annotated enhancers (13.99% **Fig 6H**), then at intergenic ATAC-Seq peaks (21.8%, **Fig S6F**), then all ATAC-Seq peaks (40.08% **Fig 6I**) and highest at GENCODE defined genes (49% **Fig S6G,H**). These results suggest that open chromatin (ATAC-Seq) is by far the best predictor of divergent nascent transcription and that relatively few annotated enhancers are subject to detectable divergent nascent transcription (**Fig 6J**). Interestingly H3K4me3 was enriched at many of the ATAC- Seq peaks, also evident in the IGV snapshots, which may reflect long range interactions with promoters or other regions enriched with H3K4me3 (**Fig 6A,B&S6A-C**). Taken together, these data strongly support the idea that many regions of the genome have open chromatin and modified histones. However, less than half of the regions with open acetylated chromatin were also subject to nascent transcription raising the interesting possibility that these regions may be primed for subsequent transcription when appropriate signals are received.

To explore this, a bioinformatic analysis was used to identify all filtered ATAC-Seq peaks (n = 45,289) with different combinations of SNU-Seq (> 0 for log_2_(x+1)), H3K27ac and/or H3K4me3 (> 1.25 for log_2_(x+1)) values to eliminate noise and reliably identify peaks (**Fig S7A-C**). After thresholding using the average signal in each peak, a “**+** /**-**“ annotation was used to summarise the properties at each ATAC-Seq peak (**Fig 7A**). 56.26% (25,476) of all ATAC-Seq peaks are enriched for both H3K27ac and H3K4me3. 31.39% (14,214) of all ATAC-Seq peaks with H3K27ac and H3K4me3 also have divergent nascent transcription (orange arrows in **Fig 7**) while 24.87% (11,262) are primed without a SNU-Seq signal (blue arrows in **Figs 7&S7**). Other combinations of marks and transcription are also evident (**Fig 7A**) but here the focus is on the potentially poised signals with ATAC-Seq peaks decorated with both H3K27ac and H3K4me3 but no transcription (blue arrows in **Figs 7&S7**). Examples of such regions in uninduced Hep3B cells are illustrated at an annotated promoter (*CD274*) (**Fig 7B**), Fantom5 annotated enhancers (**Fig S6D**) or three unannotated chromosomal locations (**Figs 7C&S7E**). Four of the five illustrated SNU-, 27ac+, K4+ ATAC+ peaks induce divergent nascent transcription after treatment with IFNγ, suggesting they are primed for transcription. These regions illustrated include one gene (divergent PDAT and pre-mRNA), two Fantom5 annotated enhancers on chromosome 15 (divergent eRNAs) and two unannotated regions with characteristics of enhancers (divergent nascent transcription) on Chr 6. One unannotated region on Chr 6 remains poised (cyan arrow in **Fig S7E**), suggesting this region might respond to different signals. A genome wide analysis revealed ≍ 11,262 regions of H3K27ac marked open chromatin lacking nascent transcription, 863 of which show IFNγ-dependent divergent nascent transcription (**Fig 7D**). 384 regions (44.5%) are located at gene promoters based on GENCODE annotations with the remainder upstream, downstream or within genes (**Fig 7D**). This analysis confirms the idea of a highly poised and responsive Hep3B epigenome revealed using SNU-Seq.

**Figure 7.**
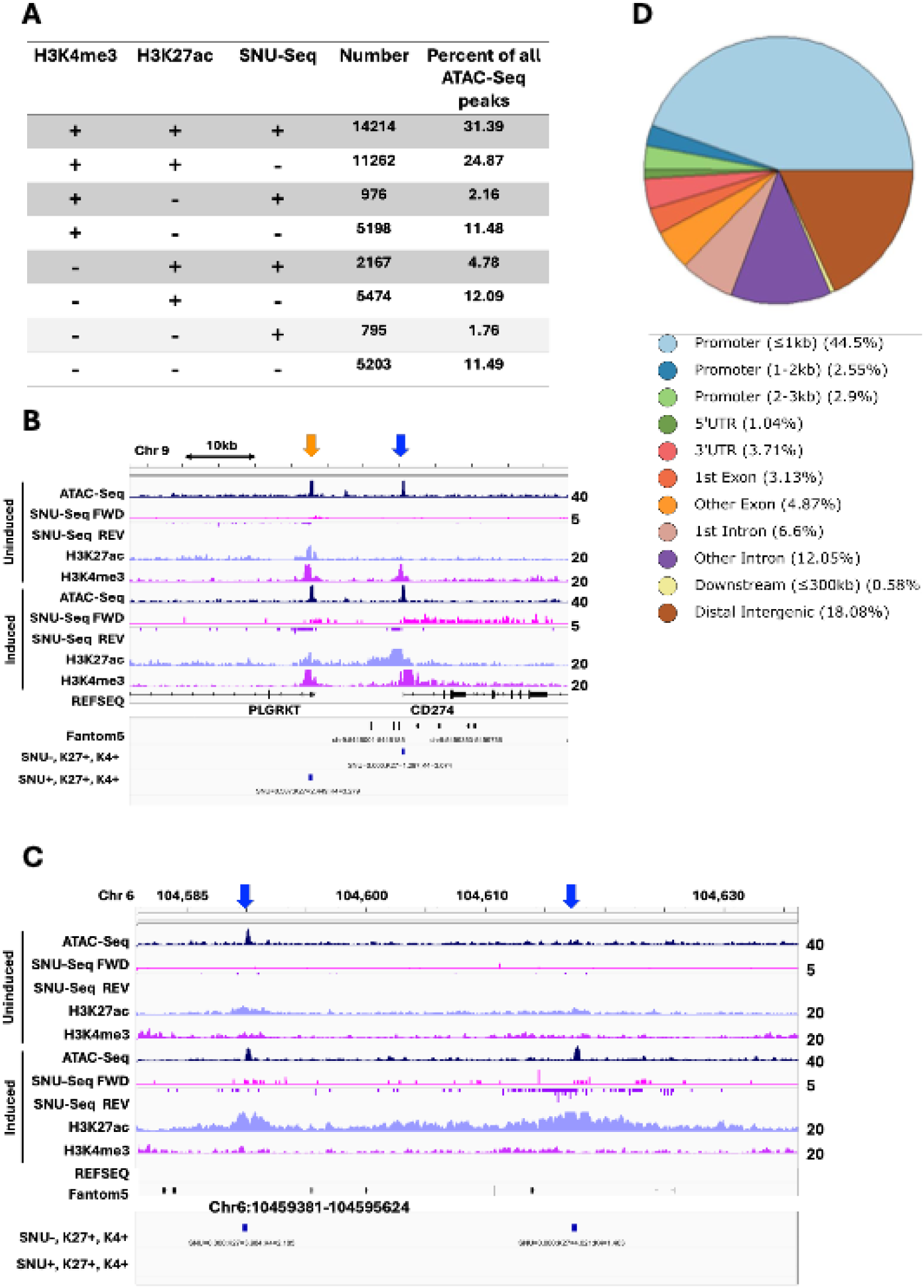
P**r**imed **sites of chromatin in Hep3B cells showing IFN**γ **inducible nascent transcription compared to regions with constitutive transcription**. **A** Number of regions of open chromatin (ATAC-Seq) and associated histone marks and nascent transcripts in Hep3B cells. **B,C** Snapshots in IGV showing chromatin features and nascent transcription around two loci selected as lacking nascent transcription until induced with IFNγ (blue arrows) or with constitutive transcription (orange arrows) at genomic regions indicated. **D** Proportion of different genomic regions with IFNγ inducible transcription. **See also Figure S7**.

## DISCUSSION

The aim of this work was to develop and validate methods to interrogate nascent transcriptomes which combine the sensitivity of the 4sU pulse-label used in TT-Seq with the single-nucleotide resolution offered by techniques such as PRO-Seq and NET-Seq, but with simple library preparation to reduce cost and improve general accessibility for routine use in mammalian cells and tissues. **S**ingle **N**ucleotide resolution 4s**U**-**Seq** (SNU-Seq) meets these criteria and allows the generation of profiles of nascent transcripts in human cells with high resolution, high sensitivity and low cost. As SNU-Seq retains full-length fragments followed by mapping the last incorporated nucleotide, there is no 5’ bias, unlike TT-Seq, where sonication or chemical fragmentation reduces the 5’ bias but does not eliminate it (16,59). In addition, unlike TT-Seq, SNU-Seq allows the presence of a promoter proximal pause (PPP) and multiple nascent polyadenylation sites to be defined in the same experiment. NET-Seq and Pro-Seq define the PPP, but not polyadenylation site usage. Finally, the single-nucleotide resolution allows accurate determination of transcription elongation rates. SNU- Seq is robust and reproducible, producing similar profiles between repeats and different human cell lines (n = 9 for HEK293 cells; n = 3 for Hep3B cells), with unique alignment of approximately 65% of initial reads. SNU-Seq also allows synthesis and decay rates to be derived, anchoring to previously obtained RNA-Seq data or, as done here, using 3’ end-Seq to sequence a small aliquot of the total RNA obtained from the same initial RNA preparation used for SNU-Seq. SNU-Seq produces an even, uniform signal over gene bodies representing productive elongation, with read density directly related to synthesis rate, and can be used directly to parametrise simulations to enable other metrics of transcription to be derived (60). In summary, unlike the other methods, SNU-Seq generates data on start site usage, site of the promoter proximal pause (PPP), transcription elongation rates and nascent polyadenylation site usage in one low-cost, straightforward, high-resolution protocol in less than two days. In addition, divergent eRNAs, a signature of active enhancers, can be resolved within genes or in intergenic regions using SNU-Seq, here validated at previously characterised enhancers (56) .

Controls were carried out to ensure that the “raw” SNU-Seq output can be used directly to assess levels of nascent transcription, PPP, elongation rates and polyadenylation site usage. There are minimal background reads in SNU-Seq preparations. Removal of rRNA reads enhances resolution of the metagenes features such as the PPP. No evidence for biased action of bacterial poly(A) polymerase at the 3’ end of nascent transcripts could be found. In addition, the gel purification step confirmed the broad size range of the 4sU- labelled nascent transcripts, ruling out extensive degradation. Uniquely, the SNU-Seq readout provides details of nascent polyadenylation during the labelling window. A control library prepared with 4sU labelled RNA but no bPAP treatment can easily be used to confirm this, and is exemplified here for *DDIT4*, *SRSF3* and *MYC*. The signal at the PAS in the SNU- Seq readout overlaps with that from the total RNA 3’end-Seq library and in the 4sU-labelled no bPAP readout, representing hPAP action during the 4sU labelling window. That these are polyadenylation sites is confirmed by the lack of signal in the control libraries for histone genes, which undergo non-hPAP-dependent 3’ end processing (61). Interestingly, SNU-Seq reports precise positioning of the transcription termination site at the histone genes, 6-12 nt downstream of the consensus ACCCA cut site after the stem loop region (62). Many other protein-coding genes, such as *HNRNPU* illustrated here, have complex patterns of polyadenylation site usage, which can be resolved by nuclear/cytoplasmic fractionation of processed transcripts (63). Certain isoforms of the transcripts are preferentially enriched in the nucleus, suggesting either selective nuclear retention or cytoplasmic turnover. Peaks in the SNU-Seq read-out match precisely with these isoforms. This confirms the very rich patterns in the use of polyadenylation sites and a great deal of potential in the way mRNAs with different 3’UTRs or with excluded exons may influence the proteome. Where multiple polyadenylation sites are used and associated with distinct outcomes in terms of levels and localisation, this appears to be hard-wired into nascent transcription, raising interesting questions as to how particular transcripts are chosen to be retained, exported or translated (64,65). SNU-Seq offers a useful tool for exploring this potential. This control was also useful to detect nascent polyadenylation at the promoter proximal region due to early termination of transcription. Although metagenes and heatmaps revealed that this is not a common event, at certain genes, such as *ZNRF3*, endogenous hPAP polyadenylation is evident close to the promoter in the sense (pre-mRNA) and antisense direction (PDATs) (25,46). The data from size fractionated 4sU-Seq confirmed that the promoter proximal pause (PPP) presents a nascent end of a ∼63 nt transcript with the 5’ end that maps to the annotated TSS or to the TSS mapped by TT-Seq, with NELF enrichment being the best predictor of the pause. In this respect, SNU-Seq shows a major advantage over TT-Seq, which is unable to resolve the PPP.

SNU-Seq has the sensitivity to detect previously unannotated non-coding nascent transcripts, including PDATs and promoter convergent antisense transcripts (PCATs) in the vicinity of promoters and divergent eRNA from regions of the genome with characteristics of enhancers. The SNU-Seq readout for non-coding nascent transcripts shows superior sensitivity to those observed in PRO-Seq, mNET-Seq and our TT-Seq readout in HEK293 cells. Although complexes such as restrictor contribute to slowing polymerase to facilitate early termination of transcription, particularly of non-coding transcripts (66), our data suggest that these transcripts extend over 1kb from the TSS, whether associated with promoters (PDATs) or enhancers (eRNAs), making them distinctly different from the short ∼63 nt transcripts with their nascent ends within RNA polymerase II at the PPP. In conclusion, 4sU- based approaches such as SNU-Seq seem particularly suitable when interested in detecting non-coding and antisense transcription. This holds true for 4sU-based TSS annotations obtained through size fractionated 4sU-Seq, too, given the high proportion of unknown TSSs that are antisense to the TSSs of already annotated protein coding genes.

We explored the potential of SNU-Seq to discover new nascent transcripts in Hep3B cells, using a chromatin analysis to define where these transcripts appear on the epigenome. This revealed several distinctive features including the presence of H3K4me3 at enhancers (67) as well as promoters, and extensive priming of the epigenome for transcription. In Hep3B cells there were over eleven thousand regions of open chromatin, defined using ATAC-Seq, flanked with H3K27ac and H3K4me3 modified nucleosomes but without nascent transcription. Rapid onset of divergent nascent transcription occurred at over 800 of these sites in response to IFNγ stimulation. SNU-Seq will be instrumental in understanding how regions of the genome with open chromatin, but no nascent transcription, respond to new signalling pathways and mapping changes in long-range interactions involving these regions, which correlate strongly with phenotype (68). Furthermore, in a similar way to the approach used in yeast (5), size fractionation of nascent transcripts and sequencing from the smallest to the largest fractions will provide base pair resolution data on the extent of contiguous transcription between overlapping genes and transcription units, which are excluded from this analysis to ensure clarity. Overlapping transcription and transcriptional interference are significant features of yeast genomes (69,70), and this approach will allow precise analysis of human genomes, particularly when responding to extracellular signals.

## METHODS

### Resources

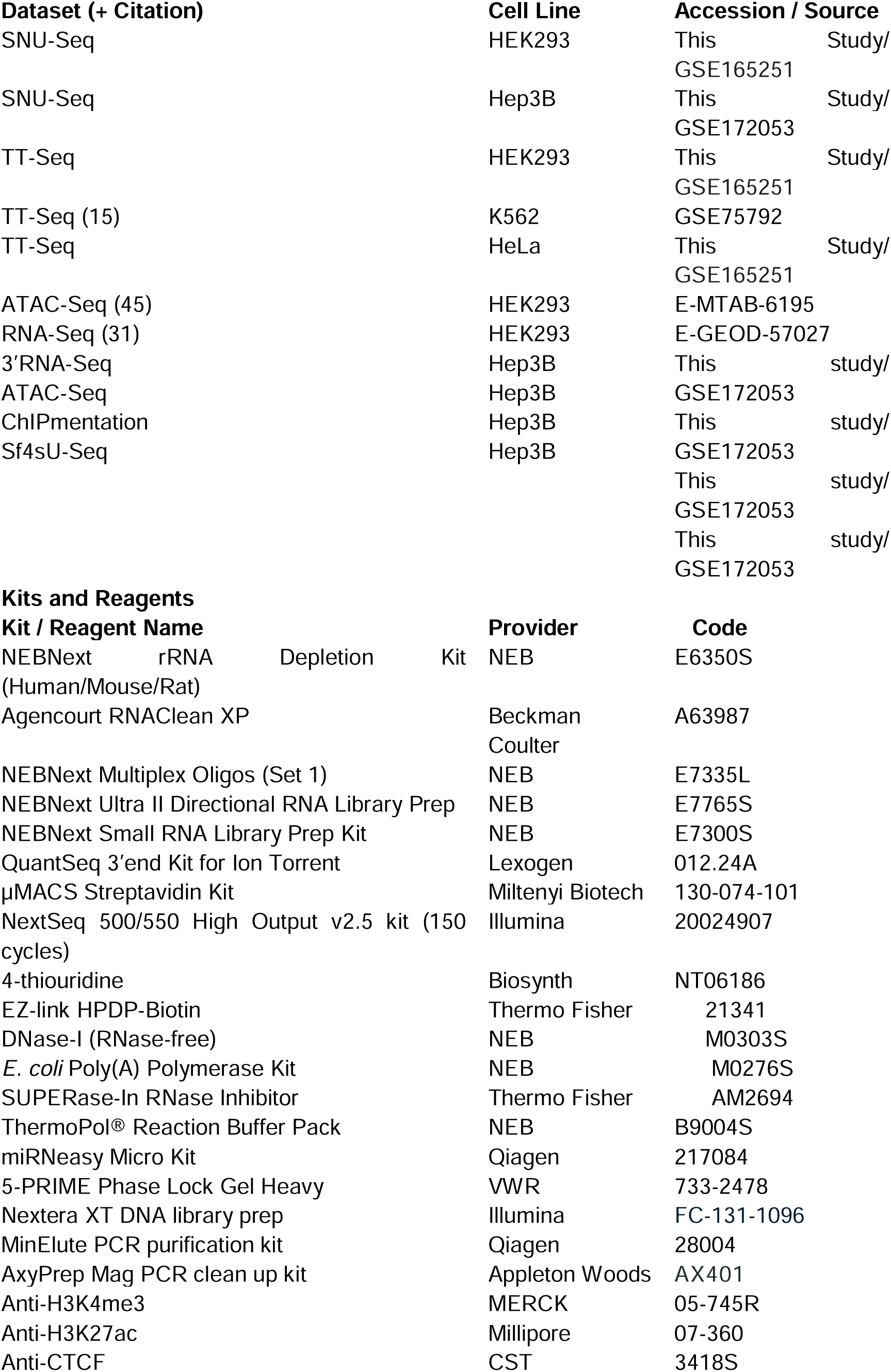
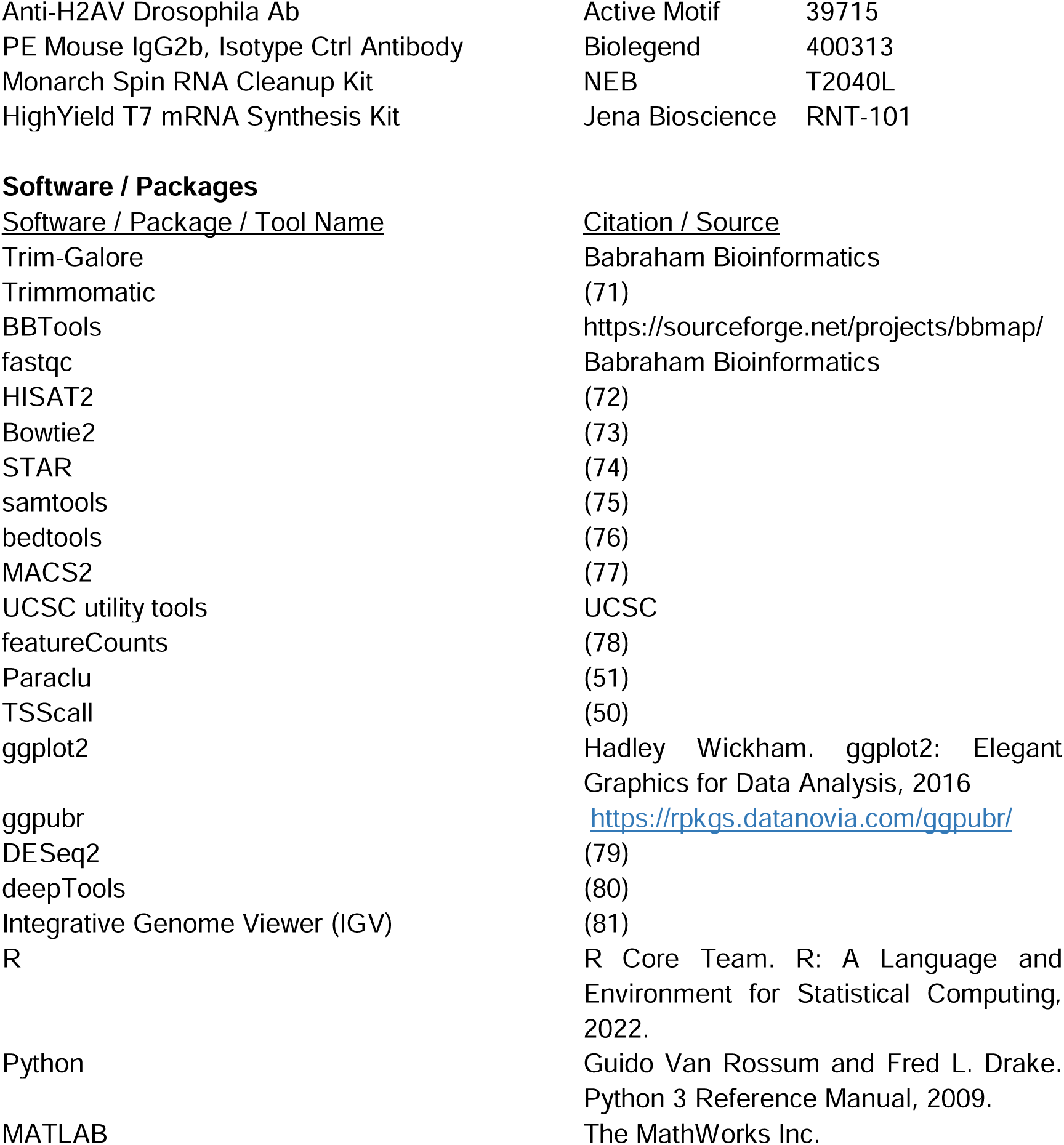

### Cell Culture

HEK293 and Hep3B cells were cultured in DMEM, supplemented with 10% (v/v) FBS and 1% (v/v) Penicillin-Streptomycin (Sigma cat #P0781). The incubator was set to 37 °C at 5% CO_2_. HEK293 and Hep3B cells were grown in three 15 cm dishes up to ∼80% confluency (∼6 x 10^7^ cells). Cells were counted on a Nexcelom Biosciences Auto 2000 Cell Counter. Hep3B cells were passaged 48hrs before harvesting at 80% confluency. After passage they were left for 24hrs to adapt, then left untreated or treated with 10 ng/ml of IFN for 0.5, 2 or 24 hrs unless stated otherwise.

### TT-Seq

TT-Seq was performed as described in the original publication (15). Sequencing libraries were prepared using the NEBNext rRNA Depletion kit for Human Cells, followed by the NEBNext II Ultra Directional RNA Library prep kit, following both kit instructions. Library quality was assessed using the Agilent Bioanalyzer with a DNA-High Sensitivity chip. Pooled libraries were sequenced on the Illumina NextSeq 500 platform using the High-Output Illumina NextSeq 500 kit. For calibrated TT-Seq with spike-ins, instructions for thio-labelled spike-in usage described in (15) were followed or prepared as follows. Briefly, 4sU-labelled spike-in mRNA encoding *Renilla* luciferase was *in vitro* transcribed (IVT) using a HighYield T7 mRNA Synthesis Kit (Jena Bioscience RNT-101). Plasmid DNA was linearized with BspQI (NEB R0712) and purified by phenol-chloroform extraction followed by ethanol precipitation to generate a template suitable for run-off transcription (1 mg template DNA used in 20 μL total IVT reaction volume). mRNA was transcribed according to the manufacturer’s instructions except from using 1.25 μL UTP (100 mM) and 0.25 μL 4sUTP (100mM, Jena Bioscience NU-1156L) to label the mRNA with 20% 4sU. The IVT reaction was incubated for 2 hr 20 mins at 37°C before treating with 2 μL DNase I (NEB M0303) in a total volume of 50 μL for 15 mins at 37°C. IVT mRNA was purified using a Monarch Spin RNA Cleanup Kit (50 μg, NEB T2040L) and eluted in 50 μL H_2_O. Concentration of purified mRNA was determined using a NanoDrop 2000 Spectrophotometer (Thermo Scientific). Integrity of IVT mRNA was confirmed by agarose gel electrophoresis and mRNA was stored at -25°C.

### TT-Seq - Sequencing Analysis

Paired-end sequencing reads were quality-checked with FastQC and then quality trimmed using Trim-Galore for a Q score below 20. Trimmed reads were then aligned to the human genome (hg38) with HISAT2 (82) using the –no-mixed and the -no-discordant flags. Aligned files in the sam format were then filtered by using samtools (75) with the flags -q 40, -f 99, - F3852, -bS. Calibration of samples (if necessary) was achieved by calculating scaling factors of spike-in counts between samples based on counts tables generated by featureCounts (78). Bedgraph and bigwig files were generated from bam files using bedtools (76), and wigToBigWig (UCSC utility tools), respectively.

### SNU-Seq

SNU-Seq libraries were generated by following the TT-Seq protocol but omitting the sonication step. Following treatments, cells were washed with cold PBS before lysing in QIAzol on ice. Samples were spiked with 4sU-labelled *Saccharomyces cerevisiae* or *Renilla* luciferase RNA (0.01%). 300 µg RNA was biotinylated. Before library preparation, nascent RNAs were polyadenylated using the NEB *E. coli* poly(A) polymerase (bPAP) following the kit instructions. The reaction with 150 ng thio-labelled RNA was left for 45 minutes at 37 °C before isopropanol precipitation. The pellet was resuspended in 11-22 µL RNase-free water. Qualities and amounts were checked on the Qubit fluorometer and the Agilent Bioanalyzer (RNA Pico Chip) or the Agilent TapeStation (RNA High Sensitivity ScreenTape).

Libraries were prepared by using the maximum RNA input amount (5 µL) following the Quant-Seq Lexogen 3’ mRNA kit (Ion Torrent or FWD for Illumina samples) instructions with 13 PCR cycles. Library qualities were checked using the Agilent Bioanalyzer (DNA High Sensitivity Chip). Following the manufacturer’s instructions, the chips prepared by the Ion Chef were then sequenced on an Ion Proton Sequencing platform or by Lexogen GmbH’s (Vienna, Austria) services on an Illumina NextSeq2000 platform. The samples were single-end sequenced with 100 nt read length (SR100). Different read depths were tried, such as ∼5 M for a batch and ∼50 M for another, as a comparison. Of note, both read depths end up with similar results, with 50 M having more duplicates but also capturing more non-coding transcripts than 5 M, indicating an ideal read depth around 20-30M.

### SNU-Seq – Sequencing Analysis

After quality control with fastqc, fastq files were trimmed with trimmomatic (71) or bbduk.sh (BBTools) to remove reads with a quality score below 20 in a sliding window of 5 bp. The poly(A) reads were also removed. Sequences were then aligned with HISAT2 with the same settings as for TT-Seq, or with STAR using “--outSAMtype BAM SortedByCoordinate -- outSAMunmapped Within --outSAMattributes All --outFilterMultimapNmax 10 -- winAnchorMultimapNmax 50 --alignSJoverhangMin 8 --alignSJDBoverhangMin 1 -- outFilterMismatchNmax 10 --outFilterMismatchNoverReadLmax 1 --outMultimapperOrder Random --alignEndsType EndToEnd --alignIntronMin 11 --alignIntronMax 0 -- alignMatesGapMax 0”. Sorted bam files were generated using samtools. For alignment, hg38 (GrCh38.p14) was used together with GENCODE (v46) annotations. Genomic A stretches were masked by using the criteria: all regions with ≥ 4 A within 6 nt but no C or T residues downstream or ≥ 12 A within 18 nt downstream of a given position, and all regions with ≥ 15 A within 18 nt upstream of a given position (24), resulted in masking ∼2.8% of the hg38 genome, including scaffolds, alternate loci and assembly patches. These masked regions, together with the blacklist regions published by ENCODE (83) were then removed from each bam file via bedtools intersect -v function. Outliers were removed by determining the top and bottom 0.5% of signal. Calibration of samples (if necessary) was achieved by calculating scaling factors of spike-in counts between samples via DESeq2, based on counts tables generated by featureCounts. Bedgraph and bigwig files were generated from bam files using bedtools, and wigToBigWig, respectively. 3’ end single nucleotide coverage was achieved by using the bedtools genomecov function’s “-3” option. A further normalisation for no bPAP bedgraph and bigwig files were done by using 3’ UTR counts obtained using bedtools map function’s “-o sum” option.

### Size fractionated 4sU-Seq

Nascent, thio-labelled RNA was generated as described for SNU-Seq. The nascent RNA was then run on a 3.5 % TBE-Urea gel (8M urea). For this, 250 ng of nascent RNA was mixed with 2x loading dye and boiled for 2 minutes at 80 °C. The gel was kept on ice and pre-run for 10 minutes at 80 V, using ice-cold ultra-pure TBE (1x) as the running buffer. Wells of the gel were washed with TBE before loading samples. Samples were then run at 80 V for 90 minutes. The gel was transferred onto a square petri dish and incubated in 1x ultra-pure TBE with SYBR Gold (1/10,000 v/v) for 5 minutes. The gel was rinsed twice in 1x ultra-pure TBE before imaging.

For purification of the small nascent RNA fraction, the gel was incubated at -80 °C for 10 minutes. The small band of 50-100 bp size was cut using a razor blade. RNA was extracted from the gel slice using dialysis: The gel slice, along with 0.8 mL 10 mM Tris-HCl (pH 7), was placed into SnakeSkin dialysis membrane sealed by two Eppendorf caps. The dialysis was run for 35 minutes at 80 V. The RNA was then purified with isopropanol precipitation and eluted in 11 uL RNase-free water. Quality and size distribution of the size-selected RNA was checked on the Agilent Bioanalyzer using an RNA pico chip.

The size fractionated nascent RNA was treated with 5’ Pyrophosphohydrolase (NEB) in ThermoPol® reaction buffer following the manufacturer’s instructions. The reaction was stopped with 1 uL 500 mM EDTA and heat-inactivated by incubation at 65°C for 5 minutes. RNA was again purified with isopropanol precipitation and resuspended in 6 uL RNase-free water. Libraries were generated using the NEBNext Small RNA Library kit for Illumina following the kit instructions. Library quality was assessed with the Agilent Bioanalyzer using a DNA High Sensitivity chip. Sequencing was performed on an Illumina NextSeq 500 platform. After sequencing, Fastq files were quality-trimmed with Trim-Galore and aligned with bowtie2 (73). Sorted bam files were generated from aligned sam files using samtools with a filtering step added to only retain reads that have a mapping quality score above 30 (- q 30). 3’end single basepair resolution was achieved by using the bedtools -bg -3 option when generating bedgraph files.

### TSS annotations

To generate TSS annotations from the size fractionated 4sU samples, 5’ end coverage was generated using the -bg -5 option (instead of -3) when generating bedgraphs from bam files in bedtools. To generate 5’end reads that only occur in nucleosome-depleted regions, ATAC-Seq data in HEK293 cells (45)(E-MTAB-6195) were used. The ATAC-Seq fastq files were trimmed with Trim-Galore, aligned with bowtie2, and converted to bam and bed files using samtools and bedtools, respectively. Subsequently, MACS2 was used to call peaks (no model assumed). The intersect option in bedtools was then used to retain only those 5’end from sf4sU-Seq signals that are located within ATAC-Seq peaks. The resulting bedgraph files were then used as an input to identify TSS cluster centres using Paraclu (51) which identifies clusters in a sliding window, and a cluster value threshold of 30 was applied. Cluster-centres were calculated using the mean position of the cluster, and TSS candidates were verified in Matlab as follows: Only those TSS candidates were assigned as true active TSSs where the summed TT-seq signal (using HEK WT 10 minute-labelled TT-Seq) in the 1000 bp downstream of the candidate TSS was at least 5 times greater than in the 1000 bp upstream of the candidate TSS. TSScall, a python code (50), was then used to identify annotated and un-annotated (novel) TSSs using the comprehensive Gencode (v29) annotation file which yielded 2955 previously annotated TSSs and 1428 novel transcription start sites. The read threshold for this was set to 5, which corresponded to an FDR of 0.001.

### Annotation Preparation for Metagene Analysis and Mathematical Modelling

GENCODE comprehensive gene annotations (n = 70,611) were filtered to keep genes within chr1 to chr22 to exclude the biased/unmapped regions in the reference genome (n = 59,950). Then, only the genes with a minimum 3.5 kb distance to each other, regardless of the strand, were kept (n = 12,224). From this subset, only the protein-coding genes ≥ 1 kb were retained (n = 2,807). Finally, the UTRs within ±100 bp of a gene end with a transcript support level 1 to 5 were detected, and they are merged if there are multiple PAS/UTRs for a gene. Only the genes with a 3’ UTR annotation were kept (n = 2,394). This is the GENCODE annotation subset that is used for metagene analysis and mathematical modelling.

### Mathematical Modelling and Determination of Transcription Constants

Synthesis and decay rates were determined based on a previously published modelling approach (15,32). HEK293 total or 3’ end RNA-Seq counts from (31) or data from this study were used. The pausing index was calculated as described (84). We used the ratio of normalised TSS (-50 nt to +200 nt) counts to gene body (TSS + 200 nt to the 3’ UTR start site) counts.

### Metagene and Data Analysis

deepTools was used for metagene calculation and visualisations. “scale-regions” mode was used to compute metagene matrices with 5 kb gene body length, 2 kb flanking regions, 20-nt bin size by averaging and skipping missing or zero-valued data. Each strand was calculated separately with strand-specific annotations, then merged with “computeMatrixOperations rbind” before visualisation. “computeMatrixOperations subset” was used to subset the matrix for the libraries of interest.

p-values were determined using the non-parametric Wilcoxon rank sum test, and the Bonferroni correction for multiple testing was applied when required. For correlations, Pearson (r) was used for the correlation between repeats based on the counts, while Kendall (τ) was used for the correlation between synthesis rates and counts, and Spearman (ρ) was used in all other cases. Principal Component Analysis (PCA) and data visualisation were performed via deepTools (multiBigwigSummary followed by plotPCA) or in R using the PCAtools and ggplot2 with ggpubr packages, respectively.

To compare the splicing levels in SNU-Seq and total RNA libraries, SPLICE-q (34) was used to calculate splicing efficiency (SE) from the bam files. The SE score refers to the number of splicing events based on the ratio of spliced to unspliced reads found around the splice junctions. The SE score (between 0 and 1) indicates a higher number of splicing events closer to 1. The mean SE was computed in each library.

To investigate the effect of the background signal (no 4sU) and the host polyadenylation signal during the pulse labelling window (no bPAP), a signal subtraction was performed for each nucleotide in the genome. For this exhaustive subtraction, deepTools bigwigCompare was utilised with “--operation subtract --binSize 1” parameters.

### Nuclear and Cytoplasmic RNA Extraction

Extraction of RNA from nuclear and cytoplasmic subcellular fractions was carried out with HEK293 cells for 3 biological replicates (65). QuantSeq 3’ mRNA-Seq Library Prep Kit for Ion Torrent (Lexogen) was applied for nuclear (500 ng input) and cytoplasmic RNA (1,700 ng input) using 13 PCR cycles. Reads were aligned to the genome build using the Ion Torrent Server TMAP aligner with default alignment settings (-tmap mapall stage1 map4). Human poly(A) site (PAS) annotations were obtained from PolyA_DB3 (85). Each PAS was extended 20 nt 3’ and 200 nt 5’ from the site of cleavage, and those that overlapped on the same strand after extension were combined into a single PAS annotation. Mapped reads were narrowed to their 3’ most nucleotide and those which overlapped with the extended PAS annotations were counted. PASs associated with non-coding RNAs and genes not in the RefSeq (86) gene database were excluded. Genes with only one PAS were also excluded.

### Chromatin analysis in Hep3B cells

ATAC-Seq was performed as previously described (87). 5 x 10^6^ Hep3B cells were washed in cold PBS and resuspended in lysis buffer (10 mM Tris-HCl pH 7.4, 10 mM NaCl, 3 mM MgCl_2_, 0.1% IGEPAL). The nuclei were pelleted and resuspended in 1X TD with 2.5 l TDE1 (Nextera XT DNA library prep kit, Illumina). These were incubated for 30 min at 37 ℃. The tagmented DNA was purified using the MinElute PCR purification kit (Qiagen, 28004). Each sample was amplified using the Nextera XT DNA library prep kit and Nextera XT index kit (Illumina) with the following thermocycler programme: hold at 72 ℃ (5 min), 98 ℃ (30 s), 9 cycles of 98 ℃ (10 s), 63 ℃ (30 s), 72 ℃ (30 s) with a final 72 ℃ extension for 1 min. The libraries were purified by adding 1.8 x volume of RT AxyPrep beads (AxyPrep Mag PCR clean up kit) to each reaction. They were then size selected with a ratio of 0.6:1 beads, to remove fragments larger than 600 bp. Libraries were run on the Agilent Bioanalyzer with a DNA-High Sensitivity chip to assess quality. Paired-end sequencing was performed on a NextSeq 500 with the 75 cycles NextSeq 500/550 High Output v2 kit (Illumina). FastQCs were performed on each dataset to ensure good quality of the run. The adapter sequences were trimmed and paired using Trimmomatic, removing reads with a quality score below 20 in a 5 bp sliding window and reads below a minimum length of 30 bp (71). Trimmed reads were aligned to the hg38 genome if un-spiked, or to a combined hg38-dm6 genome if spiked, using Bowtie2 (73). samtools were used to remove duplicates and filter out reads with a MAPQ quality score < 30 as well as mitochondrial reads (75). MACS2 was used to call peaks with a minimum FDR (q-value) of 0.01 (77). Following this, ENCODE blacklisted regions were removed (83). For visualisation of tracts, reads were normalised based on sequencing depth. Filtered BAM files were converted to bedgraphs using bedtools (76).

The ChIPmentation protocol used is largely based on the protocol published by Schmidl et al. (88). ∼10^6^ cells were washed and collected in cold PBS. The cell pellet was resuspended in room temperature PBS and fixed for 5 min with 1% formaldehyde before quenching with

0.125 M Glycine. 5000 fixed Drosophila sg4 cells (0.05%) were added as a spike-in control. The cll pellet was obtained, washed in cold PBS and incubated on ice in cold swelling buffer (10 mM Tris-HCl pH 8, 10 mM NaCl, 0.2% NP-40, 1 mM AEBSF and 1X complete mini EDTA-free proteasome inhibitor, Roche) for 10 min. The subsequent nuclear pellet was resuspended in cold lysis buffer (10 mM Tris-HCl pH 8, 1% NP-40, 0.5% Na-deoxycholate, 0.1% SDS, 1 mM AEBSF and 1X complete mini EDTA-free proteasome inhibitor, Roche) and sonicated for 90 min, 30 s on and 30 s off, at 4 ℃ (Bioruptor, Diagenode). One tenth of the sample was taken as input control. The remaining sample was incubated overnight, rotating at 4 ℃ with the antibody. Scaling factors were calculated based on spike-in counts between samples, or based on sequencing depth for CTCF ChIP, and used when converting the filtered BAM files to bedgraphs using bedtools (76). Bedgraphs were further converted to BigWig files using bedGraphToBigWig (UCSC utility tools). Differential analysis between two datasets was performed on unnormalised data using a custom R script employing the DESeq2 package (79). To identify differential peaks rather than differential gene expression, peak files for each repeat/sample were merged into a single file in which reads were counted using featureCounts and differentially assessed (78). A cut-off threshold of FDR < 0.05 was used to define statistical significance.

### Data Access

Data from the Hep3B cells is available at GSE172053 https://www.ncbi.nlm.nih.gov/geo/query/acc.cgi?acc=GSE172053.

Data on HEK293 cells is available at GSE165251 https://www.ncbi.nlm.nih.gov/geo/query/acc.cgi?acc=GSE165251

## Acknowledgements

This work was supported by: The Wellcome Trust (WT089156MA to J.M.); the BBSRC (BB/P00296X/1 to J.M., BB/Y005848/1, BB/N001184/1 to A.F.); the Leverhulme Trust (RPG- 2016-405 to J.M.); a Wellcome Trust Strategic Award (091911); BBSRC and EPSRC studentships (BB/M011224/1 to P.L. and EP/F500394/1 to T.B.); a Wellcome Trust studentship (209897/Z/17/Z to A.L.) and a Royal Society University Research Fellowship (UF120327 to A.A.). Umut Gerlevik received a PhD fellowship from the European Union’s Horizon 2020 Research and Innovation programme under the Marie Skłodowska-Curie Actions Innovative Training Network (MSCA ITN) Cell2Cell grant agreement number 860675. We thank David Aitken for preliminary SNU-Seq data.

## Author Contributions

SNU-Seq was developed and performed by PL, UG, SX and A.W. SNU-Seq in Hep3B cells together with chromatin analysis and bioinformatics was carried out by AL and UG. Bioinformatics analysis was done by PL, UG, AL, SM and CG. Mathematical modelling and determination of transcription rate constants was carried out by UG, ASK and AA. HF conducted the polyadenylation analysis supported by AF. TB, UG and PL performed the k-means clustering and downstream analysis. Data was visualized by PL, AL and UG. JM, PL, AL and UG conceived the experiments. The manuscript was written by JM with additional input from UG, PL, AA, HF, AL and other authors.

---

**Supplemental Figure S1.**
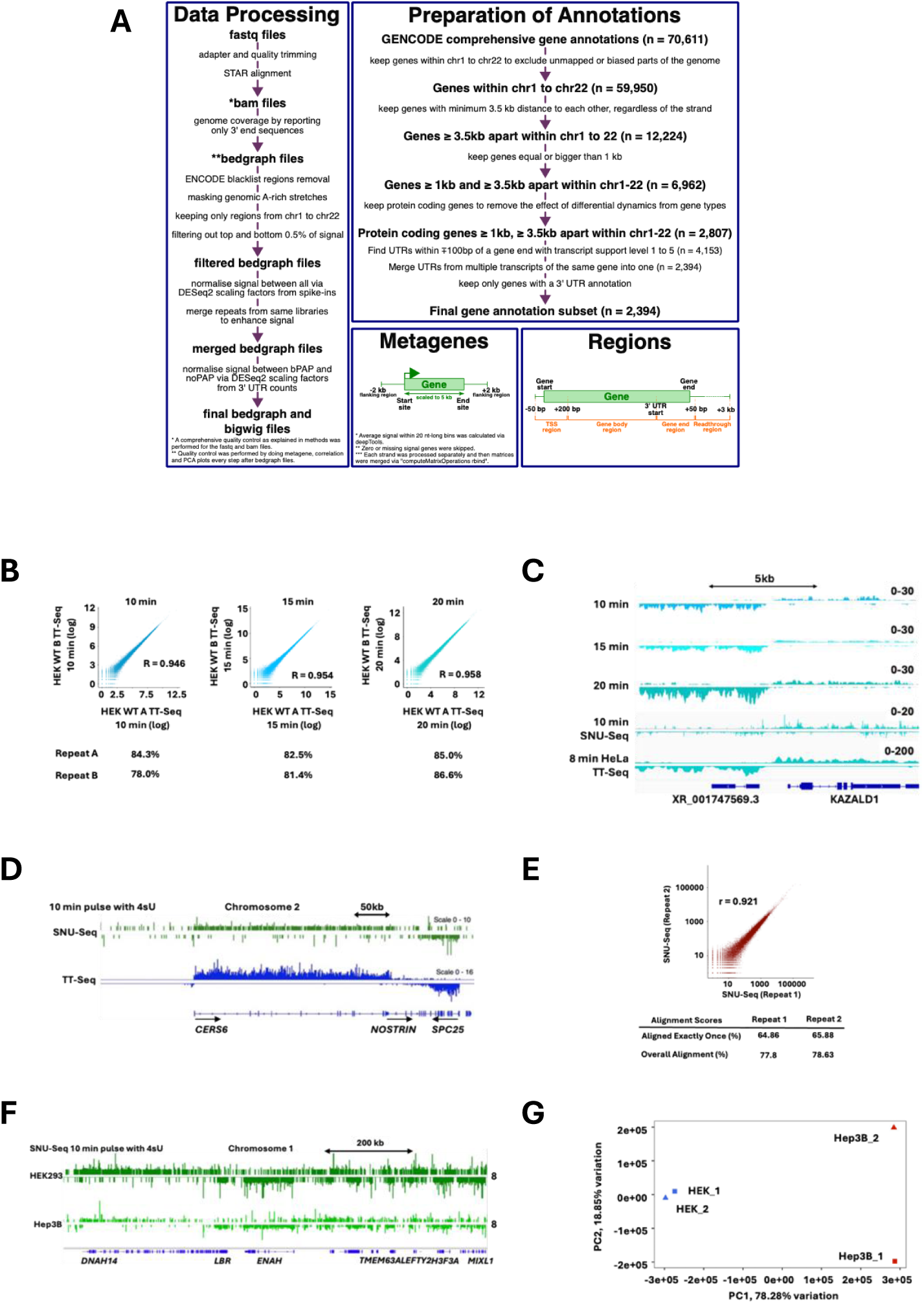
Establishment of TT-Seq and SNU-Seq in HEK293 and Hep3B cells. **A** Schematic of data processing steps. **B** Scatter plot of log-transformed read counts and alignment percentages for two repeats of 10, 15 and 20 min pulse-labelled TT-Seq experiment in HEK293 cells. **C** IGV Genome Browser screenshot of TT-Seq and SNU-Seq output at the *KAZALD1* locus from HEK293 cells and TT-Seq in HeLa cells (8 min pulse-labelled). **D** IGV Genome Browser screenshot of SNU-Seq and TT-Seq output in HEK293 cells (this study) on a region of chromosome 2. **E** Scatter plot and alignment score of SNU- Seq repeats read counts (log scale) in HEK293 cells. The Pearson correlation coefficient is shown (r = 0.921). **F** IGV screenshot displaying a comparison of SNU-Seq in HEK293 cells and Hep3B cells (this study) on a region of chromosome 1. **G** Principal component analysis of SNU-Seq in HEK293 and Hep3B cells with two repeats each. The first (x-axis) and second (y-axis) principal components are plotted against each other in a biplot. (Relates to **Figure 1**).

**Supplemental Figure S2.**
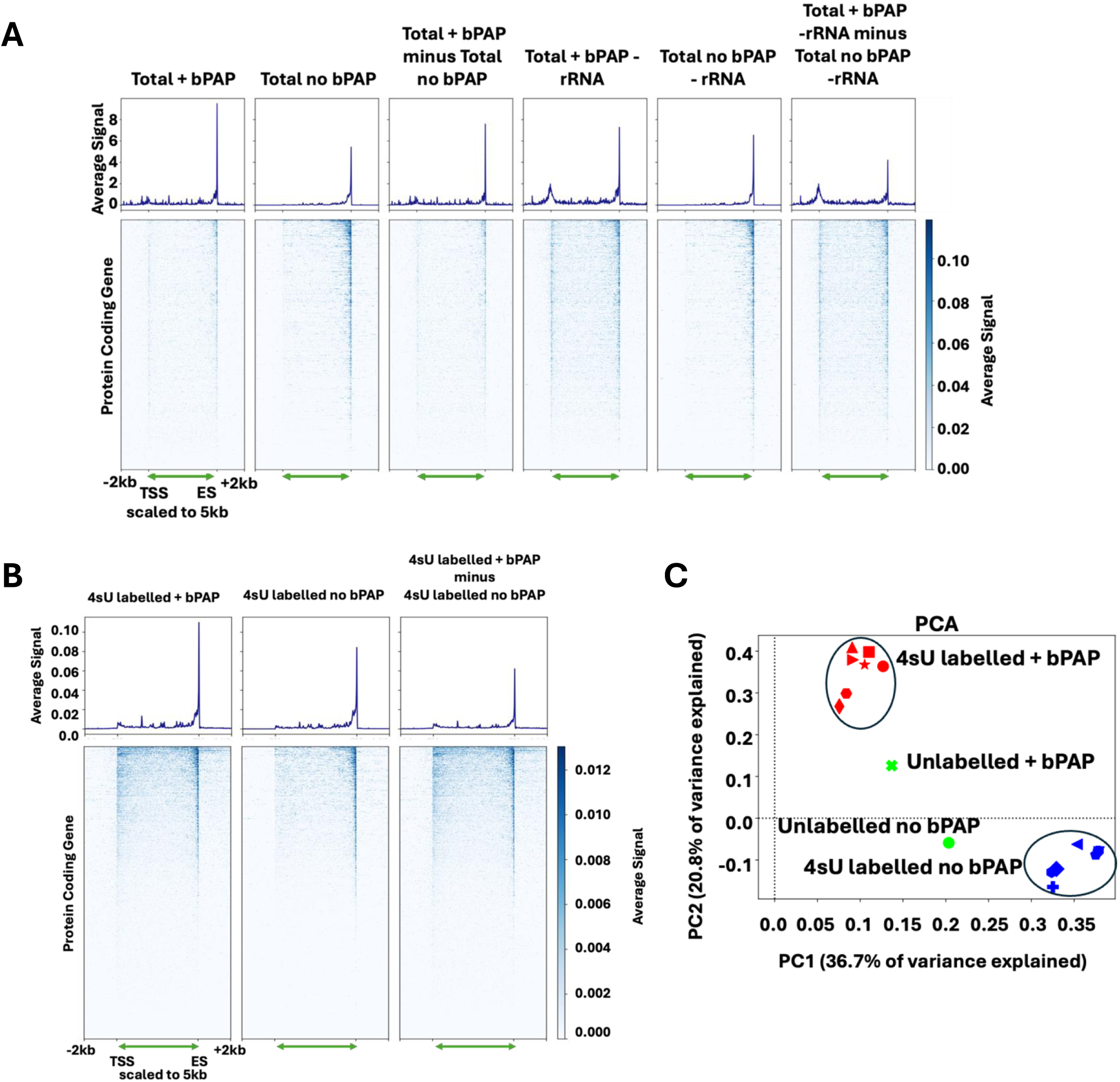
**Single-nucleotide 4sU-Seq (SNU-Seq) reports nascent transcription at high resolution**. **A,B** Metagene profiles and heat maps for 3,833 GENCODE protein coding genes after blacklisting with above threshold signal (input n = 8,743 genes) between the TSS and ES with the transcribed region scaled to 5 kb (green double headed arrow) and the flanking 2 kb up and downstream shown as log_2_ average normalised reads for total RNA (**A**), treated with or without bPAP and before or after depletion of rRNA processed as indicated. **B** Unlabelled or 4sU labelled RNA treated with or without bPAP. Samples were processed as indicated. **C** Principal component analysis of SNU-Seq in HEK293 with seven repeats for 4sU labelled samples or unlabelled samples treated with or without bPAP as indicated. The first (x-axis) and second (y-axis) principal components are plotted against each other in a biplot (Relates to **Figure 1**).

**Figure S3.**
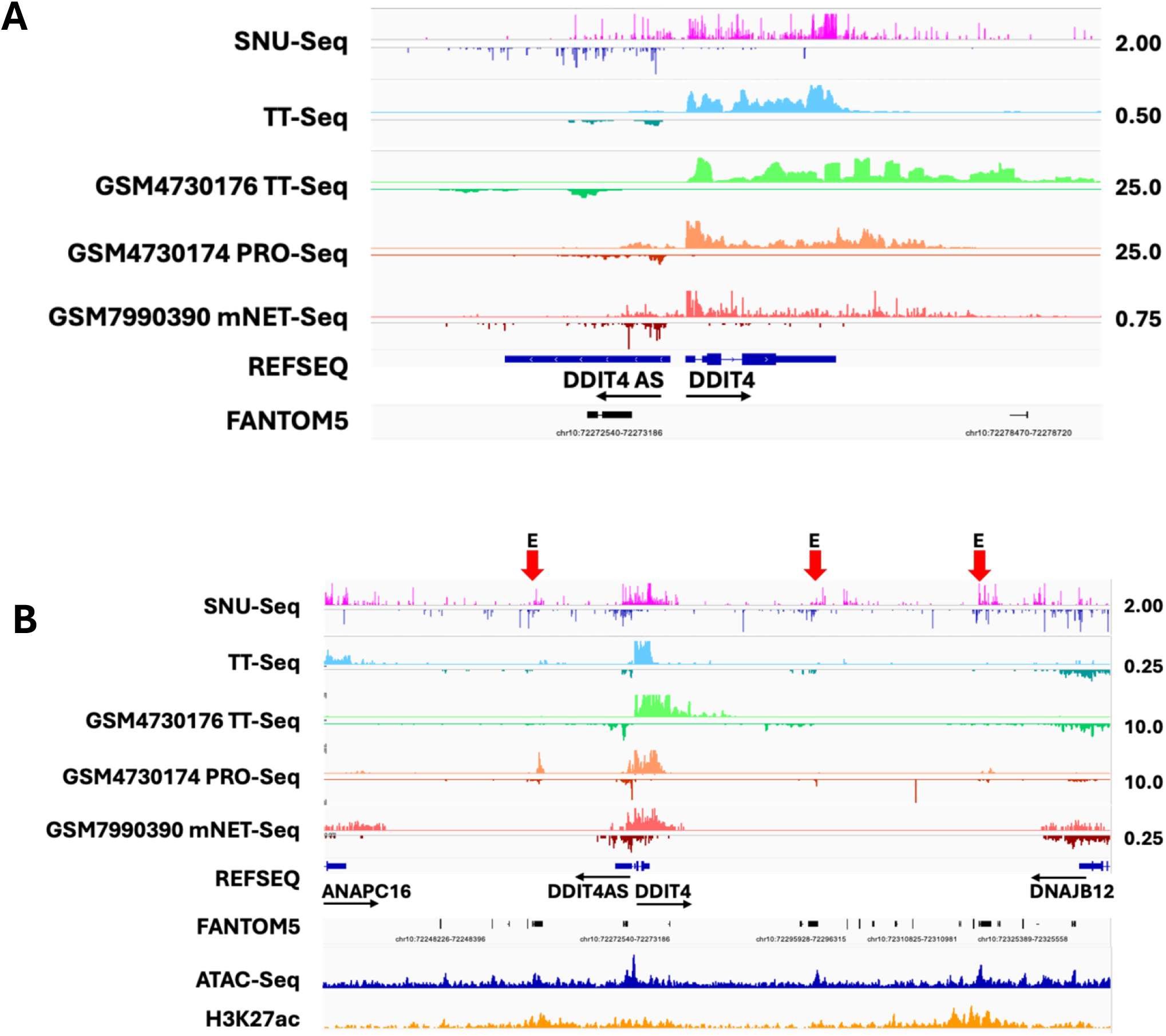
**Comparison of SNU-Seq and other methods for assessing nascent transcription. A,B**. IGV screenshot at *DDIT4* (**A**) and the region around *ANAPC16* and *DNAJB12* including *DDIT4* (**B**) displaying a comparison of SNU-Seq (10 min 4sU pulse in HEK293, this study n = 3), TT-Seq (10 min 4sU pulse in HEK293, this study n = 2), TT-Seq, PRO-Seq and mNET-Seq also in HEK293 cells from sources indicated. ATAC-Seq and H3K27ac signals are also indicated in **B** together with FANTOM5 annotations. Putative enhancers are marked with red arrows and E in **B**. Scales are indicated and for TT-Seq, PRO-Seq and mNET-Seq are adjusted in **B** to improve the detection of lower abundance eRNA transcripts compared to SNU-Seq. (**Relates to Figure 2**).

**Supplemental Figure 4.**
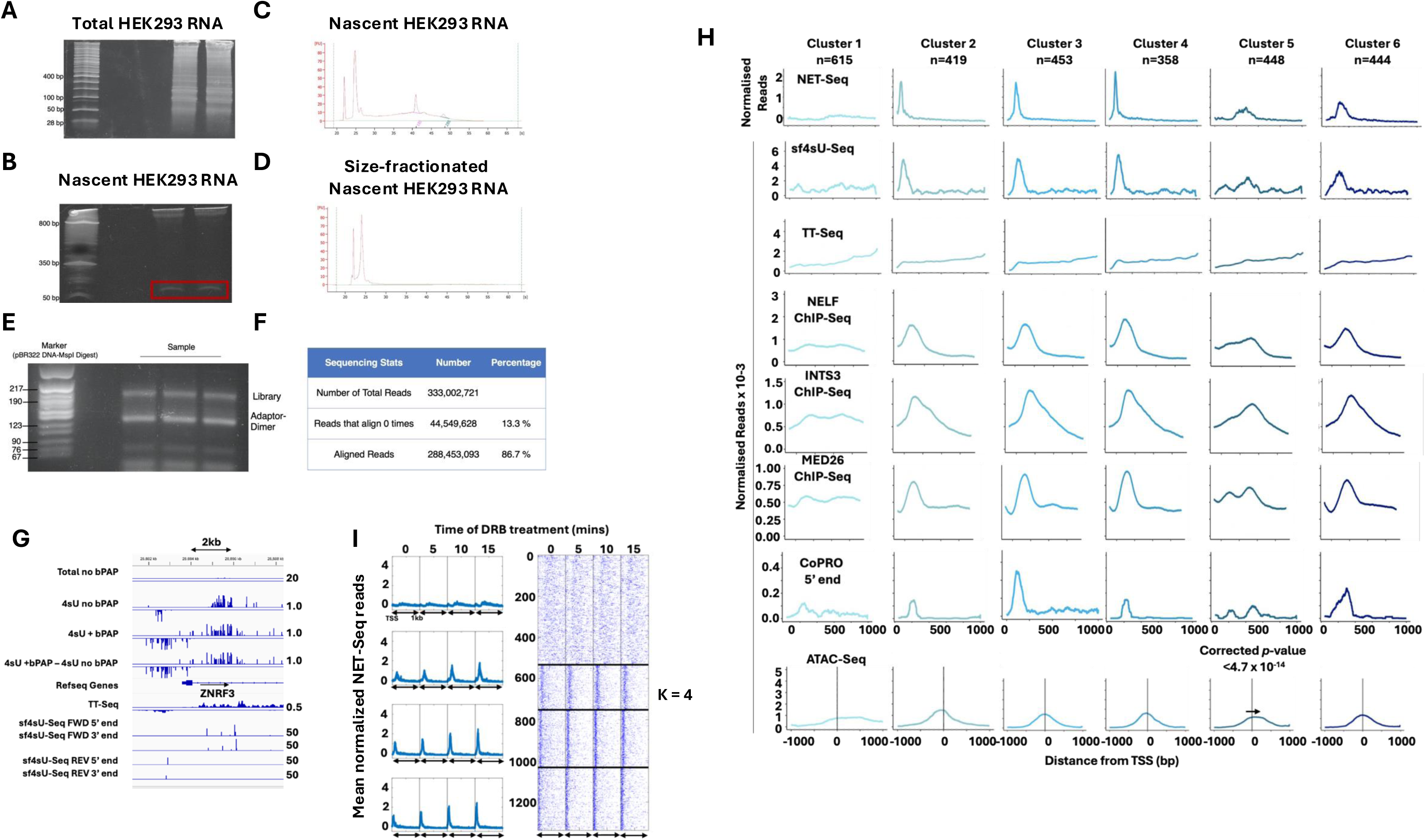
S**i**ze **fractionated 4sU-Seq captures the promoter-proximal pause**. **A-B** 3.5% TBE-urea gels run with total (**A**) and nascent 4sU labelled (**B**) HEK293 RNA. **C-D** RNA pico-chip Bioanalyzer traces of nascent HEK293 RNA before (**C**) and after (**D**) gel-based size selection. **E** sf4sU-Seq library preparation. DNA gel of sf4sU-Seq library products after adaptor ligation and PCR amplification. The top bands represent the desired library product, whereas the second-highest bands are a result of adaptor dimerisation. **F** sf4sU- Seq sequencing depth and alignment percentage. **G** Snapshot of reads around *ZNRF3* illustrating early termination and polyadenylation of promoter proximal pre-mRNA and divergent PDAT transcripts and sf4sU-Seq outputs. Scales are shown for comparison. **H** Clusters 1 to 6 based on shape of HeLa cell mNET-Seq data in **Fig. 4F**. The same cluster indices for each gene were applied to NELF, INTS3, MED26 ChIP-Seq data (HeLa cells), to TT-Seq (HEK293), to ATAC-Seq (HEK293 cells) (45) and CoPRO (K562 cells) (44) into the respective clusters. Metagenes are normalised to make every gene contribute equally. **I** Heatmaps and metagenes showing four clusters of normalised HeLa NET-Seq reads before and after treatment with the CDK9-inhibitor 5,6-dichloro-1-beta-D-ribofuranosylbenzimidazole (DRB), resulting in accumulation of RNA polymerase at the 5’ end of genes (89) for the time indicated to illustrate the response of non-pausy (cluster 1) and pausy (clusters 2-4) genes. Relates to **Figure 4**.

**Supplemental Figure 5:**
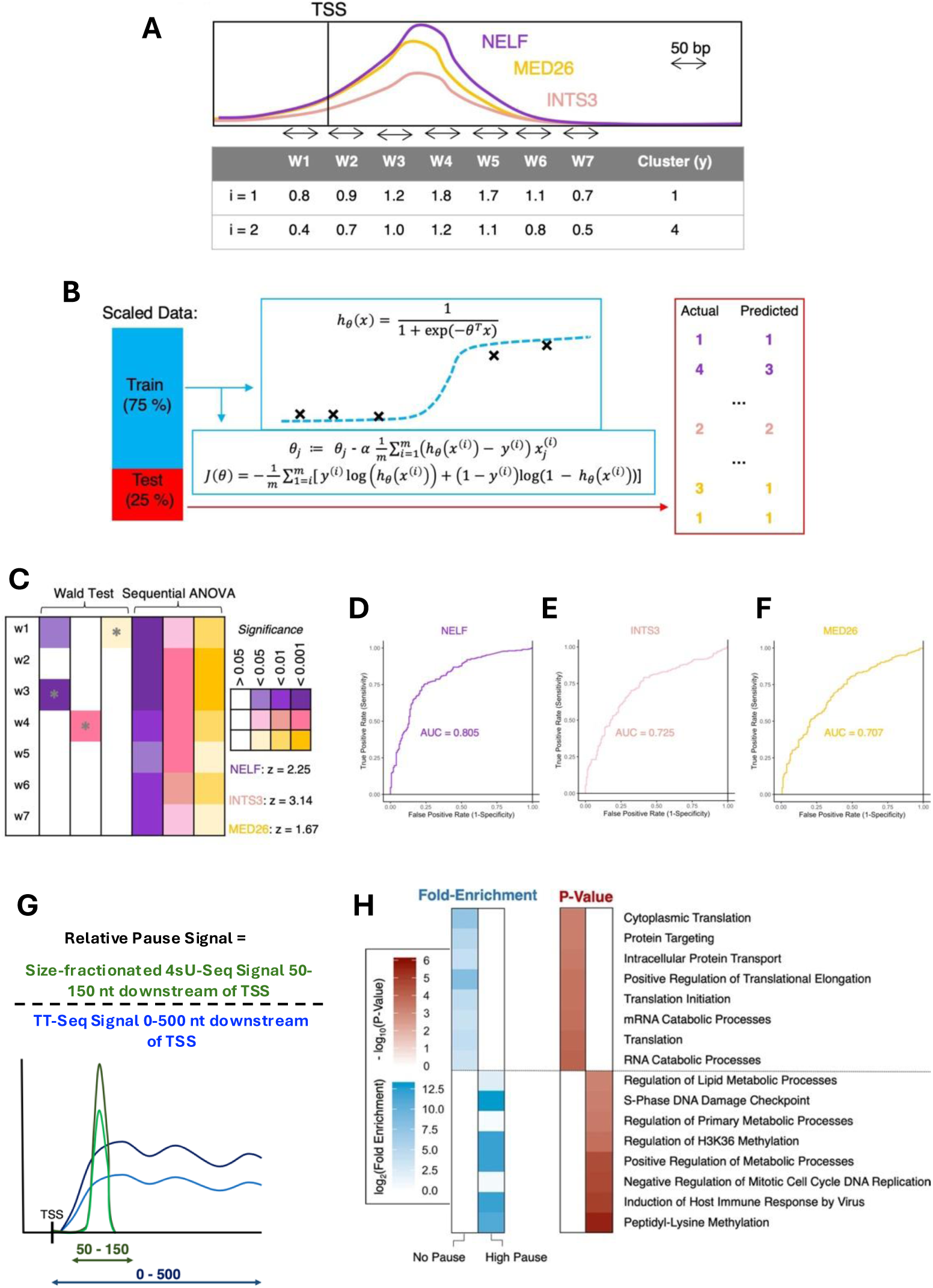
sf4sU-Seq defines the promoter proximal pause/site of early termination. **A** Classification approach of NELF, integrator, and mediator levels into NET-Seq-based clusters. ChIP-Seq data preparation for a machine learning approach. 50-bp long summed windows are generated with one window upstream (W1) and 6 windows downstream of the TSS of each gene (W2 to W7). **B** Logistic regression approach depicting the data partitioning into training and testing sets and the sigmoid hypothesis function for classification. The gradient descent and cost function equations are presented in the lower blue box. m denotes the total number of training examples (total number of genes, i.e. 1845). J denotes the cost function, i denotes the training example (i.e. the gene), and j denotes the feature (i.e. the window). The vector containing the 7 parameters is termed θ. x^(i)^ is a vector containing the summed ChIP-Seq level for the i-th gene in the 7 windows. y^(i)^ denotes the value to be predicted, i.e. the cluster. h is the hypothesis function. **C** P-values associated with the probability of parameters being significantly different from zero are displayed for NELF (purple), INTS3 (pink), and MED26 (gold), as determined by the Wald test, which tests the hypothesis whether the variable assigned to the window can be removed without affecting the prediction model. P-values in the left half of the table are determined by ANOVA, which sequentially compares the model containing only the previous variables to the model containing the respective variable. Z-values indicate the magnitude of how this parameter influences cluster classification towards clusters 2-4. A negative number would indicate that higher levels in this window would make it more likely to classify the gene as belonging to cluster 1. The window to which the maximum z-values belong is indicated with a grey star. **D-F** Sensitivity vs FPR curves and the area under the curve for NELF (**D**), INTS3 (**E**) and MED26 (**F**). **G** Calculation of the relative level of the pause signal. **H** Fold-enrichment and p-values of gene ontologies as determined by GORILLA are shown for genes that exhibit high levels of calibrated pausing (top decile) or no pausing (bottom decile). log_2_- transformed values for the fold-enrichment and negative log10-transformed values for p-values are shown. The top and bottom deciles each contained 934 genes (total number of genes was 9336). Genes with zero TT-Seq or zero SNU-Seq reads were ignored. Relates to Figure 4.

**Supplementary Figure 6.**
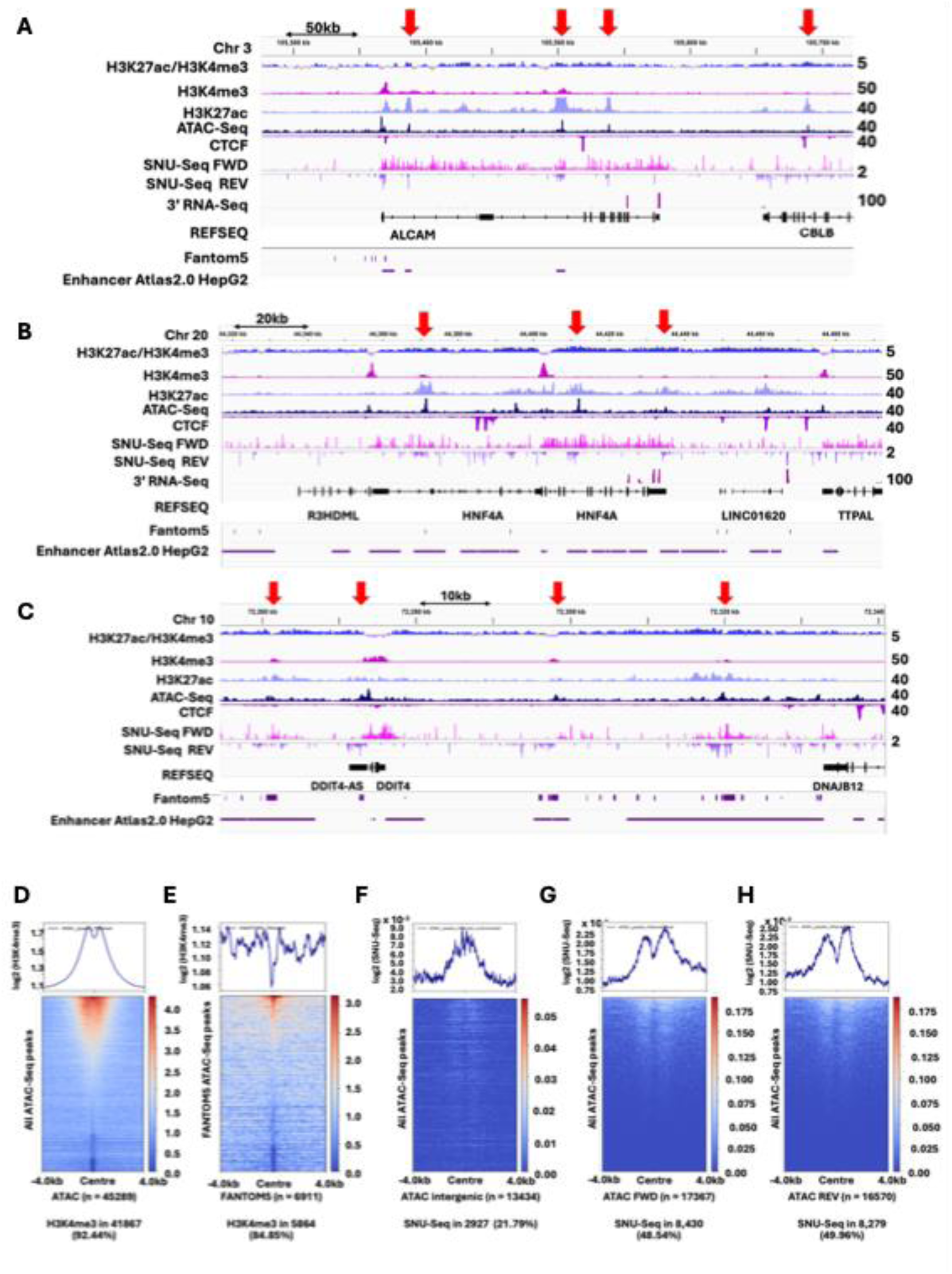
The chromatin environment and nascent transcription in Hep3B cells. A-C. Snapshots in IGV showing chromatin features and nascent transcription around three loci selected to be expressed in hepatocytes with known (*ALCAM*; **A**) or putative (*HNF4A*; **B**; *DDIT4*; **C**) enhancers indicated by red arrows. **D-H** Metagenes and heatmaps showing H3K4me3 distribution around all ATAC-Seq peaks (**D**) or FANTOM5 annotations (**E**), or SNU-Seq signals (**F-H**) around intragenic ATAC-Seq peaks, at genes on the forward (FWD) or reverse (REV) strands centred around the ATAC-Seq peaks and extending 4 kb up or downstream. The proportion of total features enriched with H3K4me3 or SNU-Seq reads over zero is shown below. **Relates to Figure 6**.

**Figure S7.**
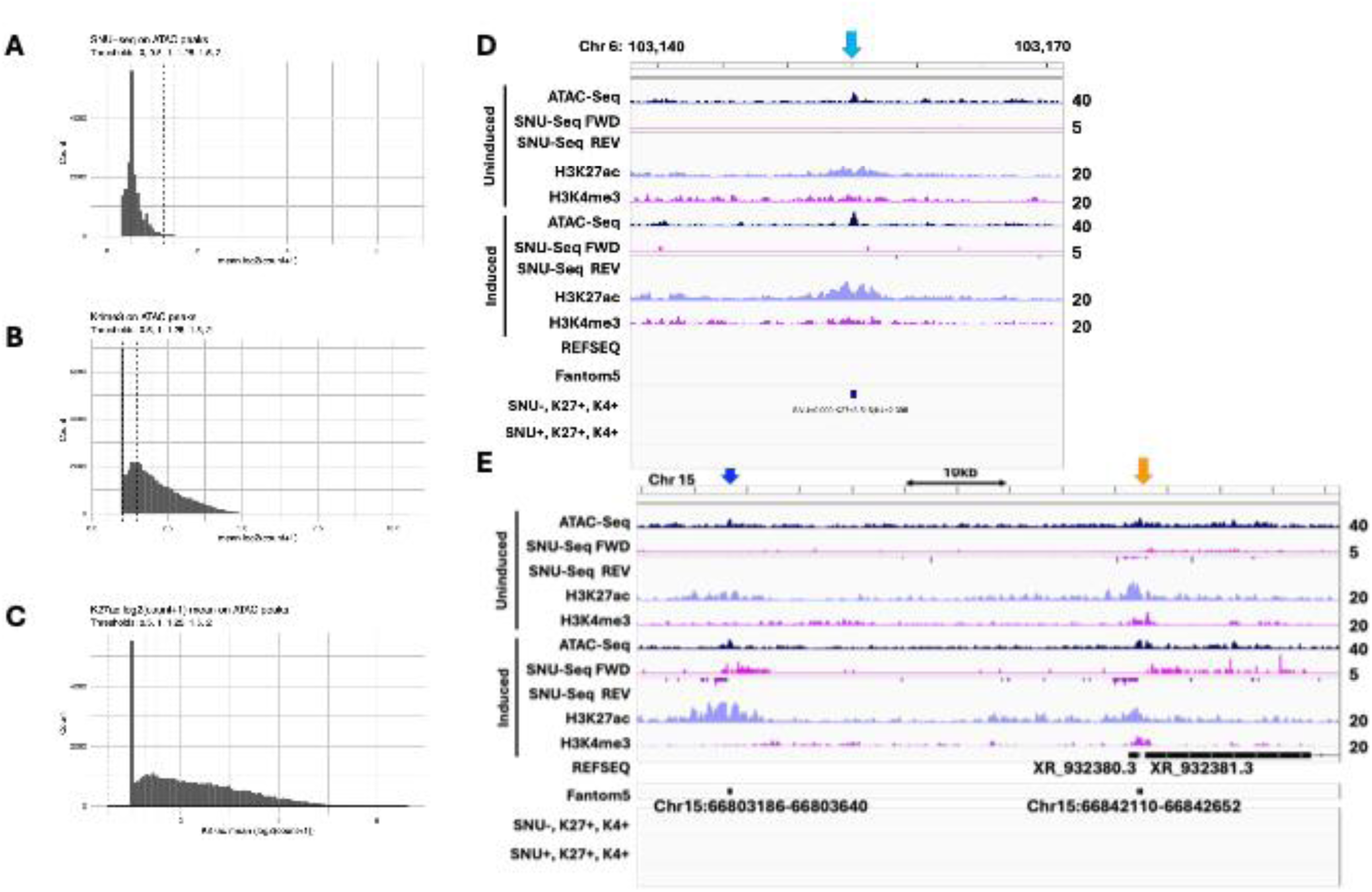
**Primed sites of chromatin in Hep3B cells showing IFN**γ **inducible nascent transcription compared to regions with constitutive transcription**. **A-C** Classifying significant levels of H3K27ac, H3K4me3 and SNU-Seq. **D,E** Snapshots in IGV showing chromatin features and nascent transcription around two loci selected as lacking nascent transcription even when induced with IFNγ (cyan arrows) or at FANTOM5 annotated enhancers with (orange arrows) or without (blue arrows) nascent transcription until induction. **Relates to Figure S7**.

**Supplemental Table 1.**
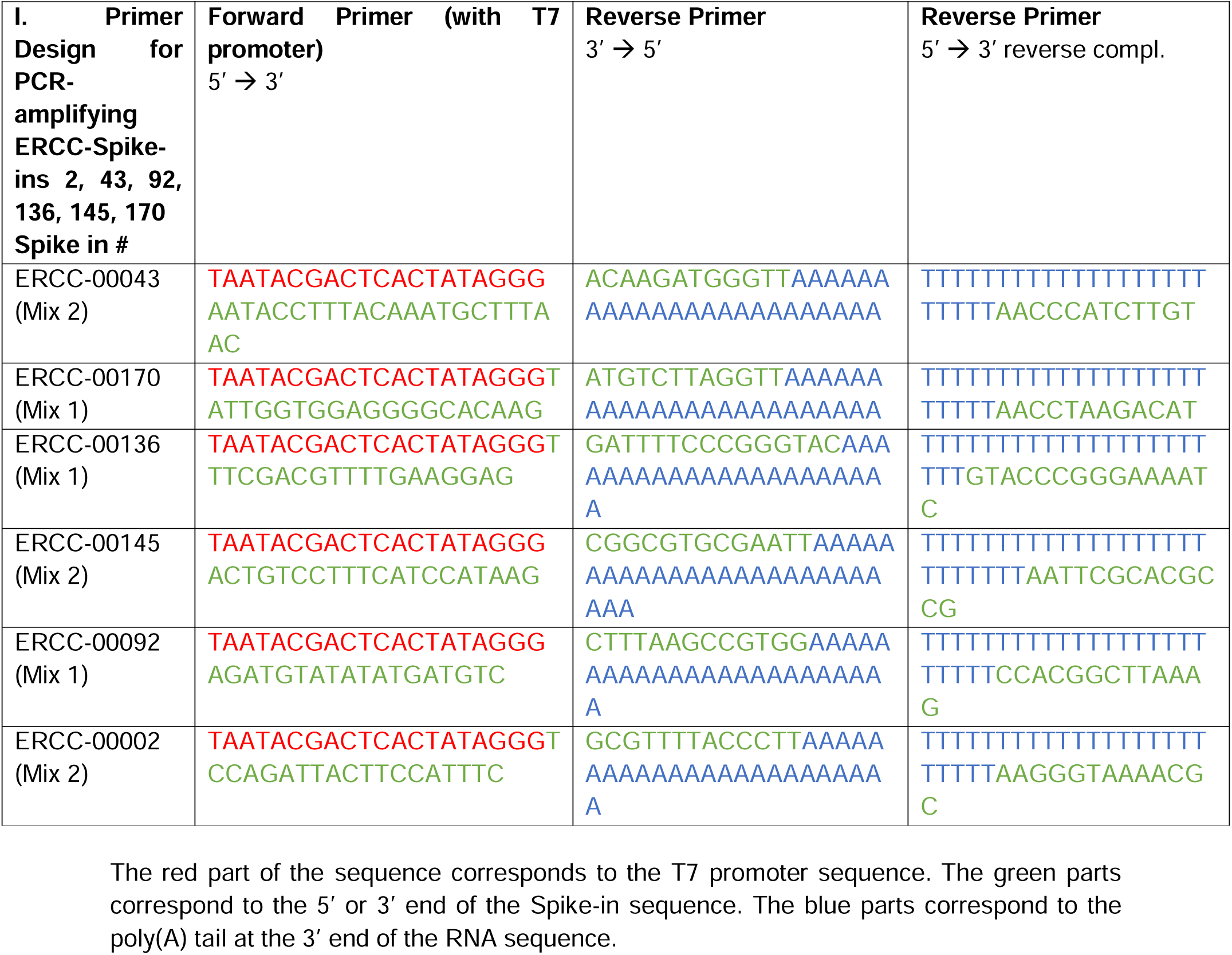
ERCC-Spike-in Mix Preparation for TT-seq.

